# Structure-informed Language Models Are Protein Designers

**DOI:** 10.1101/2023.02.03.526917

**Authors:** Zaixiang Zheng, Yifan Deng, Dongyu Xue, Yi Zhou, Fei Ye, Quanquan Gu

## Abstract

This paper demonstrates that language models are strong structure-based protein designers. We present LM-Design, a generic approach to reprogramming sequence-based protein language models (*p*LMs), that have learned massive sequential evolutionary knowledge from the universe of natural protein sequences, to acquire an immediate capability to design preferable protein sequences for given folds. We conduct a *structural surgery* on *p*LMs, where a lightweight structural adapter is implanted into *p*LMs and endows it with structural awareness. During inference, iterative refinement is performed to effectively optimize the generated protein sequences. Experiments show that LM-Design improves the state-of-the-art results by a large margin, leading to 4% to 12% accuracy gains in sequence recovery (*e*.*g*., 55.65%/56.63% on CATH 4.2/4.3 single-chain benchmarks, and *>*60% when designing protein complexes). We provide extensive and in-depth analyses, which verify that LM-Design can (1) indeed leverage both structural and sequential knowledge to accurately handle structurally non-deterministic regions, (2) benefit from scaling data and model size, and (3) generalize to other proteins (*e*.*g*., antibodies and *de novo* proteins).

## 1. Introduction

Proteins are 3D-folded linear chains of amino acids that govern biological functions such as transcription, translation, signaling, and cell cycle control. Recently, the promise of learning to understand and design proteins from data via generative deep learning has led to an ongoing paradigm shift apart from the long-established physics-based methods.

Designing protein sequences that fold into desired structures, namely structure-based protein (sequence) design, is one of the most important problems in bio-engineering. Significant progress has been made by several latest deep generative model-based approaches (Ingraham et al., 2019; Jing et al., 2020; Hsu et al., 2022; Dauparas et al., 2022; Gao et al., 2022). These approaches formulate structure-based protein design as an end-to-end graph-to-sequence learning problem, where an encoder-decoder model ℳ_*θ*_ : 𝒳 → 𝒮 is tasked with predicting protein sequence 𝒮 given a protein backbone structure 𝒳. Typically, supervised learning is performed on such models given a certain amount of protein structure-sequence pair data.

Albeit deep generative models showing revolutionized capability in this field, we argue that the current neural structure-based protein design approaches are not necessarily at their best in designing more plausible proteins as two major obstacles remain and hinder further progress:

### (i) Limited experimentally determined protein structure data

For example, the known protein structures in the commonly-used CATH (Orengo et al., 1997) dataset are multiple orders of magnitude smaller (*<* 0.1%) than the sequence data in the UniRef (Suzek et al., 2015) sequence database (**Fig. 1D-E**). As structure-based protein design is essentially a conditional sequence learning problem, the protein sequence distribution is crucial yet remains elusive for generally data-hungry generative models due to limited data. Therefore, they fail to holistically explore the protein sequence space and tend to yield sub-optimal sequence predictions for folds. Despite being partly remedied by data-augmented approach (Hsu et al., 2022), additional predicted structure data and trainable model parameters at scale demand compute and storage overheads.

**Figure 1.**
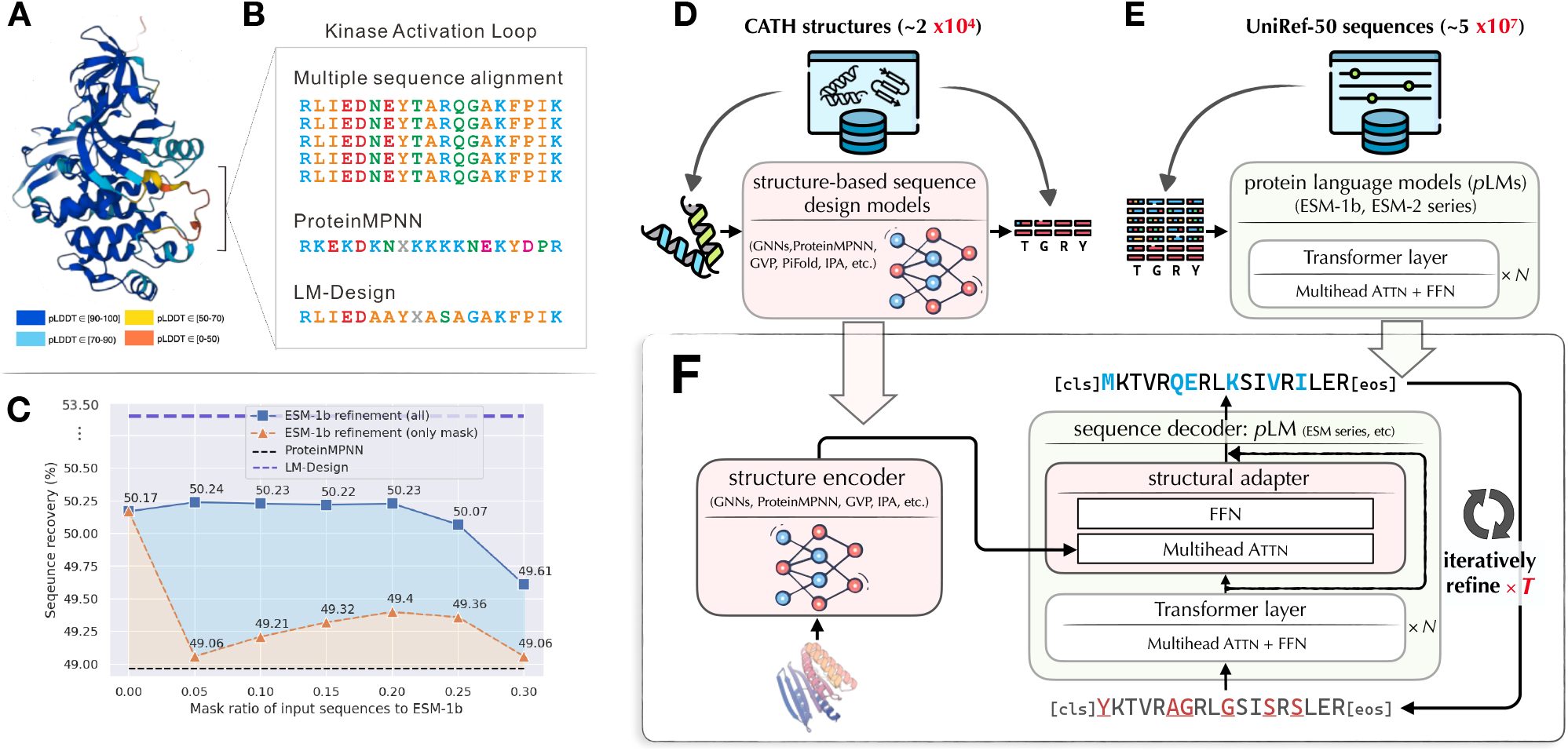
Overview. **(A)** Case study of Tyrosine kinase activation loop. Ribbon diagram shows the structure of Tyrosine kinase mapped with AlphaFold2 pLDDT score. The activation loop is characterized with low pLDDT scores suggesting flexible conformations; **(B)** Multiple sequence alignment of the activation loop showing this sequence is highly evolutionary conserved. Predictions from ProteinMPNN and LM-Design are shown; **(C)** Preliminary study on refinement ability for *p*LMs. Here ESM-1b took as input the predictions of ProteinMPNN; **(D)** Illustration of neural structure-based protein sequence design, and **(E)** protein language models; **(F)** Overall illustration of LM-Design, where **the wonderful colored protein structure image is credited to RFDiffusion** (Watson et al., 2022).

### (ii) Challenge of structurally non-deterministic regions

From a biological perspective, protein structures are sometimes not sufficiently informative, especially for those flexible regions such as loops and exposed surfaces (Towse & Daggett, 2012). In these regions, residue identities can, hypothetically, be less correlated with the structural context while sequential knowledge is way more useful yet largely neglected. We verified this hypothesis and found that existing purely structure-based approaches were prone to produce functionally invalid sequences for these regions (**Fig. 1A-B**).

Therefore, the sequential information should be better utilized for structure-based protein design. Inspired by the impressive progress of large language models (LLMs) in natural language processing (NLP) (Devlin et al., 2019; Radford et al., 2018; Brown et al., 2020), recent literature in protein research has also demonstrated the emergent evolutionary knowledge of proteins in protein language models (*p*LMs, Rives et al., 2019; Lin et al., 2022; Hu et al., 2022), learned from the universe of massive protein sequence data. Such comprehensive and thorough sequential knowledge of *p*LMs can help probe functional properties and even predict protein structures from single sequences without the need for explicit evolutionary homologs (*e*.*g*., MSAs). Thus, an exciting research question naturally arises:

Since *p*LMs are such strong sequence learners, *can we leverage p*LM*s to make better structure-based protein design?*

If so, rather than as protein sequence encoders, *p*LMs can possibly be repurposed as sequence generative models (since they are learned to reconstruct corrupted protein sequences), *prompted* by the desired structure to generate sequences, making the most of the acquired sequential evolutionary knowledge. How to best achieve this goal, however, is non-trivial and remains under-explored (we will discuss our preliminary attempts that uses *p*LMs for “post-editing” to provide insights that motivate our proposal in §2), thus deserves to be comprehensively studied.

In this paper, we show that language models with *structural surgery* are strong protein designers without using abundant training data. We propose LM-Design, a generic approach to reprogramming sequence-based protein language models (*p*LMs) to design protein sequences of a desired fold. As shown in **Fig. 1F**, we conduct a *structural surgery* on a *p*LM (*e*.*g*., ESM-1b), where a lightweight *structural adapter* is implanted to endow *p*LMs with structural awareness by access to an *arbitrary* additional structure encoder (*e*.*g*., ProteinMPNN). During inference, iterative refinement is performed to optimize the generated protein sequence until convergence when the prediction can no longer be improved.

We highlight our contributions and findings as follows:

- We introduce LM-Design, a generic approach that transforms *p*LMs to protein design models via *structural surgery*. LM-Design yields preferable protein sequences for desired structures, while being model-agnostic, modularizable, parameter- and data-efficient.
- Experiments show that LM-Design advances the state-of-the-art methods by a large margin, achieving 55.65% and 56.76% sequence recovery on CATH 4.2 and CATH 4.3 for single-chain proteins, and *>*60% for protein complexes. LM-Design can be also combined with data augmentation (Hsu et al., 2022), where additional large amounts of predicted protein structures by AlphaFold 2 (Jumper et al., 2021) are leveraged.
- In particular, we find that LM-Design can accurately handle structurally non-deterministic regions (*e*.*g*., functional loops and exposed surfaces) thanks to the learned sequence knowledge from *p*LMs, while previous methods typically fail. We also find that LMDesign can indeed be structurally sensitive, thereby better determining the nuanced sequential specificity of those protein groups of high structural similarity.
- We also show that LM-Design can synthesize diverse and structurally valid sequences. We further evaluate zero-shot generalizability of LM-Design in designing proteins of unseen categories, including antibodies and *de novo* proteins, and observe superb performance.

We highlight that the goal of this study to propose LMDesign is not to compete but instead to complement current neural structure-based sequence design models. We hope that LM-Design can become a powerful, universal, and easy-to-use tool as a “wrapper” that helps integrate the advances of both protein sequence learning (*e*.*g*., *p*LMs) and structure learning (*e*.*g*., geometric/graph NNs and protein structure prediction), facilitating future protein research.

## 2. Preliminaries

### 2.1. Structure-based Protein (Sequence) Design

Structure-based sequence design problem (*a*.*k*.*a*., protein inverse folding) is to find, given a protein backbone structure of interest, an amino acid sequence that will fold to this structure (Dauparas et al., 2022). While physics-based approaches tackle sequence design as an energy minimization problem (Dahiyat & Mayo, 1997; Street & Mayo, 1999) like the Rosetta (Alford et al., 2017), recent advances in deep learning (DL) methods have demonstrated great promise in generating plausible amino acid sequences for desired protein structures (Ingraham et al., 2019; Hsu et al., 2022).

#### Problem Formulation

Neural structure-based protein design can be formulated as an end-to-end graph-to-sequence learning problem. Formally, a parameterized encoder-decoder neural model ℳ_*θ*_ is tasked with predicting the protein sequence for a protein backbone structure,

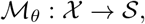

where for a protein of length *L*, 𝒮 = {*s*_*i*_ ∈ Cat(20)|1 ≤ *i* ≤ *L*} is a residue sequence of 20 types of amino acids, and 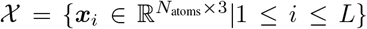 denotes the spatial coordinates in 3D space for the residues of the desired protein structure with *N*_atoms_ backbone atoms (*e*.*g*., *N, C*_*α*_ and *C*, with *O* optionally). The learning objective is to find the model parameter *θ* that maximizes the conditional log-likelihood *p*(𝒮| 𝒳; *θ*) given sufficient protein structure-sequence paired data. This enables us to design sequences of maximum likelihood, or with sampling algorithms when the diversity and novelty of designs are taken into account.

#### Overview

The general workflow of these approaches (**Fig. 1C**) is as follows: (1) A desired protein backbone structure 𝒳 is first represented as a *k*-nearest-neighbor (*k*-NN) graph in 3D space with geometric features attaching to nodes and edges of the graph; (2) A graph neural network-based encoder then takes as input the featurized graph and maps it to structural representations; and (3) Finally, a sequence decoder consumes the encoded structural representations and accordingly predicts a sequence of amino acids 𝒮 that is expected to fold into the target protein structure 𝒳, in which an autoregressive decompo-sition 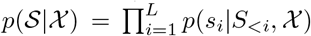 is typically applied. Notably, a recent work (Gao et al., 2022) found that a fully non-autoregressive factorization (Gu et al., 2018), *i*.*e*., 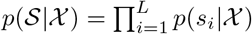 can help achieve a faster and better result. We will elaborate on the probabilistic models for structure-based protein design (see §3.1).

### 2.2. Protein Language Models

Language models (LMs) trained on large-scale sequence data have shown extraordinary advances and led to a significant paradigm shift in NLP, boosting machines in understanding human languages (BERT/MLM-style, Devlin et al., 2019) and synthesizing human-like text (GPT/CLM-style, Radford et al., 2018). Analogies between protein sequences and human languages have long been noted (Yang et al., 2019; Ferruz & Hö cker, 2022). As such, it is exciting to expect cutting-edge techniques of language modeling can tackle protein-related problems, especially for generative scenarios such as protein design.

Typically, protein language models (*p*LMs, **Fig. 1D**) approximate the protein sequence distribution *p*(𝒮) via pseudo-likelihood (Salazar et al., 2020), wherein ∏_*i*_ *p*(𝒮_*i*_| 𝒮_−*i*_) over a partially corrupted sequence (by being randomly masked, replaced or kept up to certain schedules) is maximized. Although the only training objective is to identify missing amino acids, a high success rate necessitates the model to learn intricate information within its sequential input, *e*.*g*., underlying evolutionary correlations and tertiary topology.

Please refer to Appendix §G for more detailed discussions on the related work.

### 2.3. Proof-of-Concept

As aforementioned, there are two major challenges for current neural structured-based protein design: (1) from the (conditional) sequence learning perspective, ***the lack of abundant protein structure data***, thus a model is hard to properly explore the protein sequence space through data-intensive supervised learning; and (2) from the biological perspective, ***protein structures are not necessarily always informative***, especially for those flexible regions such as loops and exposed surfaces (Towse & Daggett, 2012; Hsu et al., 2022). In these cases, residue identities are less correlated to the spatially associated tertiary structure while sequential evolutionary knowledge can be more decisive. For example, as shown in **Fig. 1A-B**, the activation loop in Tyrosine kinase has multiple conformations and is not well spatially constrained. However, it is known to play an important function in regulating the activity of the protein. When pure structure-based modes, *e*.*g*. ProteinMPNN (Dauparas et al., 2022), is used to design this functional loop, it was prone to produce functionally invalid repeated sequences.

Based on the above observations, we conjecture that protein language models (*p*LMs) can have promise for circumventing these two problems since they have learned from largescale protein sequence data, thereby protein sequential evolutionary knowledge is supposedly acquired. To verify our hypothesis, we started with a quick sanity check by studying whether a trained *p*LM has the capability to refine predicted sequences generated by currently advanced strong structure-based approaches. Specifically, we inspected the pretrained ESM-1b (Rives et al., 2019) for *p*LM together with the ProteinMPNN (Dauparas et al., 2022) for the protein design model, both are popular and openly accessible. We tested on CATH 4.2 test split, where sequence predictions of the ProteinMPNN were fed as input into the ESM-1b.

As shown in **Fig. 1C**, without any training, an immediate accuracy improvement is achieved. This promising observation boils down to two properties of *p*LMs: (1) acquired sequential evolutionary knowledge, and (2) learning to denoise from corrupted protein sequences. This sheds light on the use of *p*LMs to boost protein sequence design. Note that in this naive *post-editing* manner, *p*LMs cannot access the structure directly, hence the potential of exploiting structural information to steer *p*LMs for structure-based sequence design is not fully utilized. In the next section, we will take a further step to devise a generic framework that makes sequence-level *p*LMs and structures best integrated by the proposed *structural surgery*.

## 3. Reprogramming *p*LMs for Structure-based Protein Design with Structure Surgery

### Overview

We proposed a generic framework named LMDesign that reprograms *p*LMs to yield high-quality protein sequences from desired backbone structures. A general illustration is shown in **Fig. 1F**. To endow pLMs with structural awareness, we introduce a lightweight *structural adapter* (Houlsby et al., 2019) into *p*LMs (*e*.*g*., ESM-1b) such that the emergent ability of acquiring protein geometric information from an arbitrary structure encoder is unlocked. During inference, iterative refinement is performed to optimize the generated protein sequence until the prediction could no longer be improved.

For brevity, we will take ProteinMPNN and ESM-1b as a running example hereafter. Note that, however, the proposed framework is model-agnostic, regardless of the choices for any structure encoders and *p*LMs. It can accommodate any other strong structure-based design models, *e*.*g*., Pi-Fold (Gao et al., 2022), and ESM-2 series as *p*LM alternatives. This also implies that LM-Design can be highly modularizable, especially when pretrained parameters of the structure encoder are available. These two merits help LM-Design attain further performance bonus (see $4).

### 3.1. Training

To better accommodate *p*LMs that are tasked with masked language modeling (MLM, Devlin et al., 2019) as the learning objective, our framework is accordingly established in a denoising auto-encoding fashion. More concretely, we use conditional masked language modeling (CMLM, Ghazvininejad et al., 2019), which is proven to be very useful for conditional sequence generation in NLP (Gu et al., 2018), as the learning scheme. CMLM shares similarities with the BERT-style objective of sequence representation learning, but is more suitable for the generative purpose. Formally, given backbone structure 𝒳 and sequence 𝒮= *S*_masked_ ∪ *S*_obs_, CMLM requires the model to predict a set of target amino acids *S*_masked_, which are randomly masked, from the remaining observed residues

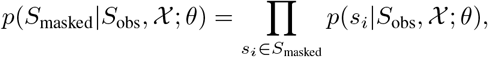

Here a conditional independence assumption over identities of target residues *s*_*i*_ ∈ *S*_masked_ is made, given 𝒳 and *S*_obs_.

We remark that such a conditional independence assumption is almost true for structure-based protein design from the viewpoint of probabilistic graphical models (PGMs), wherein graphically represented protein structure implies that each amino acid is primarily dependent on its spatially nearest neighbors rather than considerably distant ones. In our preliminary experiments, we found that existing strong models (*e*.*g*., ProteinMPNN) trained with CMLM consistently outperformed their original autoregressive formulation, where only “left” contexts rather than all neighbors get considered. This indicates that such an assumption can effectively exploit the underlying structural information, thereby better formulating structure-based protein design.

### 3.2. Inference with Iterative Refinement

To predict protein sequences from a given structure, our goal is to sample sequences 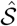 that have high likelihoods *w*.*r*.*t. p*(𝒮| 𝒳). Particularly, 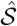 with the maximum likelihood can be obtained via greedy deterministic decoding: 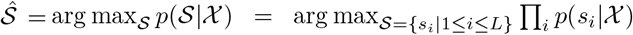 Notably, LM-Design is trained to reconstruct a protein native sequence from its corrupted version, which enables it to iteratively refine the predicted sequence in a coarse-to-fine manner towards a better one (Savinov et al., 2021).

Concretely, to sample in such an iterative refinement manner, we follow the structure of the Markov process and sample sequentially 𝒮^(*t*)^ ∼ *p*(𝒮^(*t*)^ | 𝒮^(*t*−1)^, 𝒳) by recycling the *p*LM-based decoder for some fixed number of steps *T*, starting from an initial sequence 𝒮^(0)^. Here the initial sequence 𝒮^(0)^ can be drawn from a weaker proposal distribution parameterized by a simple linear projection to amino acid vocabulary from the features of the structure encoder^1^. In our primary case where ProteinMPNN (Dauparas et al., 2022) serves as the structure encoder, we let this linear projector be its original output head, hereby 𝒮^(0)^ can be regarded as the output of the ProteinMPNN.

We can tune the number of steps *T* for a good accuracy-efficiency trade-off. Larger *T* usually leads to better prediction with high latency, whereas one-shot parallel generation, as a special case, can be achieved by setting *T* = 1 when efficiency is prioritized.

#### Temperature-based Sampling

To control the diversity and speed of convergence, we consider a modified function of the categorical distribution we want to sample from such that 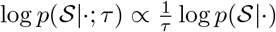, where *τ* is the temperature.

## 4. Experiments

In this section, we evaluate LM-Design on a variety of benchmarks for fixed backbone protein sequence design (including single-chain proteins and multi-chain protein complexes). We start with evaluations on designing single-chain proteins using standard CATH 4.2 and 4.3 (Orengo et al., 1997) benchmarks as in (Ingraham et al., 2019; Hsu et al., 2022). We then go for assessing capabilities of designing multi-chain complexes using the dataset curated and used in Dauparas et al. (2022). We instantiate the structure encoders in LM-Design using the same hyperparameters as the corresponding backbone structure models, and the dimensions of the structural adapter are thereby determined. Moreover, comprehensive and thorough analyses on diverse settings of our interest are conducted thereafter, including sequence accuracy *vs*. diversity, effects of scaling *w*.*r*.*t*. data and model size, structural validity, and generalizations to other proteins (*e*.*g*., antibodies and *de novo* proteins). Finally, systematic bio-informatics assessments are presented to help dive deep into the intrinsic properties of LM-Design.

Please refer to Appendix for more detailed experimental settings and additional results of all the evaluations.

### 4.1. Benchmarking Fixed Backbone Protein Design

#### 4.1.1. Single-chain Protein Design

**Tab. 1** shows the results of LM-Design in comparison to the recent strong baselines on the CATH (Orengo et al., 1997) benchmark, including the current strongest ones. We highlight our primary findings as follows:

**Table 1.** Performance comparison between LM-Design and different baseline approaches on CATH 4.2 and CATH 4.3 datasets. LMDesign’s results are obtained by 5-cycle iterative refinement with `argmax` operator (*i*.*e*., no sampling). †: benchmarked results are quoted from Gao et al. (2022). ‡: “Single-chain” in Hsu et al. (2022) is defined differently. ESM-1b 650M is used as the default *p*LM.

| | Models | Trainable/Total<br>Params. | Perplexity (↓) | | | Median Recovery (% , ↑) | | |
| --- | --- | --- | --- | --- | --- | --- | --- | --- |
|  |  |  | Short | Single-chain | All | Short | Single-chain | All |
| CATH 4.2 | †Structured Transformer (Ingraham et al., 2019) | 1.56M/1.56M | 8.39 | 8.83 | 6.63 | 28.14 | 28.46 | 35.82 |
|  | †GVP (Jing et al., 2020) | 1.0M/1.0M | 7.23 | 7.84 | 5.36 | 30.60 | 28.95 | 39.47 |
|  | †ProteinMPNN (Dauparas et al., 2022) | 1.9M/1.9M | 6.21 | 6.68 | 4.61 | 36.35 | 34.43 | 45.96 |
|  | PiFold (Gao et al., 2022) | 6.6M/6.6M | 6.04 | 6.31 | 4.55 | 39.84 | 38.53 | 51.66 |
|  | ProteinMPNN + CMLM [ProtMPNN-CMLM] | 1.9M/1.9M | 7.16 | 7.25 | 5.03 | 35.42 | 35.71 | 48.62 |
|  | LM-DESIGN (ProtMPNN-CMLM) | 6.9M/659M (1.0%) | 7.02 | 6.67 | 4.63 | 35.71 | 38.61 | 53.26 |
|  | LM-DESIGN (pretrained ProtMPNN-CMLM: *fine-tune*) | 6.9M/659M (1.0%) | 7.33 | 6.83 | 4.63 | **36.47** | **40.91** | **54.62** |
|  | LM-DESIGN (pretrained ProtMPNN-CMLM: *freeze*) | 5.0M/659M (0.7%) | **7.01** | **6.58** | **4.41** | 35.19 | 40.00 | 54.41 |
|  | PiFold (reimpl.) | 6.6M/6.6M | 6.94 | 6.82 | 4.69 | 36.67 | 38.21 | 50.22 |
|  | LM-DESIGN (PiFold) | 11.9M/664M (1.7%) | **6.77** | **6.46** | **4.52** | **37.88** | **42.47** | **55.65** |
| CATH 4.3 | GVP-large (Hsu et al., 2022) | 21M/21M | 7.68 | ‡6.12 | 6.17 | 32.60 | ‡39.40 | 39.20 |
|  | GVP-Transformer (Hsu et al., 2022) | 142M/142M | 8.18 | ‡6.33 | 6.44 | 31.30 | ‡38.50 | 38.30 |
|  | +1.2M AF2 predicted data (Hsu et al., 2022) | 142M/142M | 6.05 | ‡4.00 | 4.01 | 38.10 | ‡51.50 | 51.60 |
|  | LM-DESIGN (pretrained GVP-TransEncoder: *freeze*) | 6.3M/800M (0.8%) | 6.61 | 7.64 | **3.92** | 42.31 | 38.64 | **56.49** |
|  | ProteinMPNN + CMLM | 1.9M/1.9M | 6.31 | 6.32 | 4.85 | 40.30 | 39.02 | 48.25 |
|  | LM-DESIGN (ProtMPNN-CMLM) | 6.9M/659M (1.0%) | 5.96 | 5.85 | 4.48 | 44.12 | 44.05 | 54.05 |
|  | LM-DESIGN (pretrained ProtMPNN-CMLM: *fine-tune*) | 6.9M/659M (1.0%) | 5.94 | 5.80 | 4.20 | 45.65 | **46.15** | 55.92 |
|  | LM-DESIGN (pretrained ProtMPNN-CMLM: *freeze*) | 5.0M/659M (0.7%) | **5.88** | **5.66** | **4.19** | **45.71** | **46.15** | **56.38** |
|  | PiFold (reimpl.) | 6.6M/6.6M | 5.88 | 5.55 | 4.47 | 42.86 | 43.69 | 50.68 |
|  | LM-DESIGN (PiFold) | 11.9M/664M (1.7%) | **5.66** | **5.52** | 4.01 | **46.84** | **48.63** | **56.63** |

##### (1) LM-Design is more data-efficient and advances state-of-the-arts methods with a large margin without any additional data

On the more commonly-used CATH 4.2 benchmark, we observe that improving protein featurizers capability structure encoders, from the vanilla message-passing (graph) neural networks (Ingraham et al., 2019) to more complicated ones (Jing et al., 2020; Dauparas et al., 2022; Gao et al., 2022), indeed facilitate performance to some extents, but are limited to the under-representative protein sequence distribution due to the data shortage of experimentally determined CATH datasets, which is expected to be mitigated by *p*LMs. Our motivation has primarily been verified through our illustrative ProteinMPNN + ESM-1b setting, wherein by taking advantage of massively pretrained *p*LMs, the proposed LM-Design improves ProteinMPNN + CMLM by 4.4% on CATH 4.2 (48.62% → 52.99%) and 5.8% on CATH 4.3 (48.25% → 54.05%), setting the new state-of-the-arts without using any augmented data.

##### (2) LM-Design can be modularizable and further benefit from pretrained structure models

Instead of learning structure encoders from scratch, LM-Design can leverage pretrained structure models as encoders, which can be finetuned together with the structural adapter or kept frozen. As little difference in sequence recovery, we suggest that freezing is the best practice. In this case, only a tiny proportion of parameters (*i*.*e*., those of the structural adapter) are trainable, and LM-Design quickly converges with better results in a negligible overhead of 10 epochs. Notably, we also study LM-Design with encoder from GVP-Transformer (Hsu et al., 2022), which was built on 1.2M AF2 predicted data, giving rise to a ∼5% further improvement.

##### (3) The more advanced the structure encoder becomes, the stronger our model-agnostic LM-Design performs

Since our aim is to devise LM-Design as a general-purpose framework that can make the most of the progress of protein structure models, we study whether LM-Design can get boosted as if meets stronger structure encoders. We built a variant of LM-Design upon PiFold (Gao et al., 2022), a most recent and performant structure-based design model. As we can tell from **Tab. 1**, LM-Design improves PiFold by at least 5.43% in terms of sequence recovery, yielding impressive 55.65% and 56.63% on CATH 4.2 and 4.3 datasets. These results demonstrate that LMDesign is a general-purpose approach for structure-based sequence design, which is compatibility-friendly and hence fully leverages the advances of protein structure learners.

Combining **(2)** and **(3)**, LM-Design helps structure-based protein design make more gains with less pain, thanks to the breakthrough progress and the open-source efforts of the whole community, standing on the shoulders of giants.

#### 4.1.2. Multi-chain Protein Complex Design

A protein functions only when it docks, combines, and interacts with other macro- and macro-molecules, composing multi-chain protein complexes. As such, studying protein sequence design for multi-chain assemble structures is crucial for drug design. This incentivizes us to evaluate whether LM-Design can better handle protein complex design.

We use the multi-chain complex dataset curated by Dauparas et al. (2022) and use the same training settings as in our single-chain scenario. We summarize our results regarding protein complex design in **Tab. 2**. We first notice that CMLM (Ghazvininejad et al., 2019) can better formulate and train ProteinMPNN than the original autoregressive version with teacher-forcing. Upon a more competent system of ProteinMPNN + CMLM, LM-Design yields a near 60% sequence recovery of multi-chain protein assembles. When further integrated with *p*LMs at scale (*i*.*e*., ESM-2) or better structure encoder (*i*.*e*., GVP-TransformerEncoder), it can even achieve more impressive scores of 61.49% and 62.16%, respectively. These results show that LM-Design can not only design single-chain proteins, which are mostly studied in previous works but also be used for designing multi-chain protein complexes. This makes LM-Design more general-purpose in terms of the categories and scenarios where it can be deployed, and opens opportunities to use it for designing specific protein complexes (see §4.3), such as antigen-antibody or protein-ligand assemblies.

**Table 2.** Performance on multi-chain protein complex dataset (in median recovery). Results of the original ProteinMPNN and GVP-Transformer were obtained using publicly available checkpoints.

| Models | Rec. (↑) |
| --- | --- |
| ProteinMPNN (Dauparas et al., 2022) | 50.00 |
| ProteinMPNN + CMLM [ProtMPNN-CMLM] | 54.39 |
| LM-DESIGN (ProtMPNN-CMLM + ESM-1b 650M) | 59.10 |
| LM-DESIGN (pretrained ProtMPNN-CMLM: freeze) | **59.43** |
| LM-DESIGN (pretrained ProtMPNN-CMLM: fine-tune) | **59.43** |
| LM-DESIGN (ProtMPNN-CMLM + ESM-2 650M) | 59.81 |
| LM-DESIGN (ProtMPNN-CMLM + ESM-2 3B) | **61.49** |
| LM-DESIGN (pretrained GVP-TransEncoder: freeze + ESM-1b 650M) | **62.16** |

### 4.2. Analyses: Diving Deep into LM-Design

#### 4.2.1. Structural Validity using AlphaFold 2

Given that experimental assessment is not available, we leverage the most famous *in silico* structure prediction protocol, *i*.*e*. AlphaFold 2 (AF2, Jumper et al., 2021), to evaluate the structural validity of our designs. Here we use the pLDDT score of AF2 as the evaluation metric and follow the evaluation configurations as in Dauparas et al. (2022), where AF2 takes as input only single sequences (native sequences and our designs) while no multiple sequence alignments (MSAs) are provided. We redesign all 1120 proteins in CATH 4.2 test split. As shown in **Fig. 2, LM-D****esign****’*s redesigns are predicted, by AF2, to adopt the given backbone structures more confidently than the native sequences***, implying higher structural stability of our redesigns. We postulate this is because, in this challenging setting where no co-evolutionary information of homologous sequences is exposed to AF2, LM-Design exploits the full potential of sequential co-evolutionary knowledge that *p*LMs learns from massive protein sequences.

**Figure 2.**
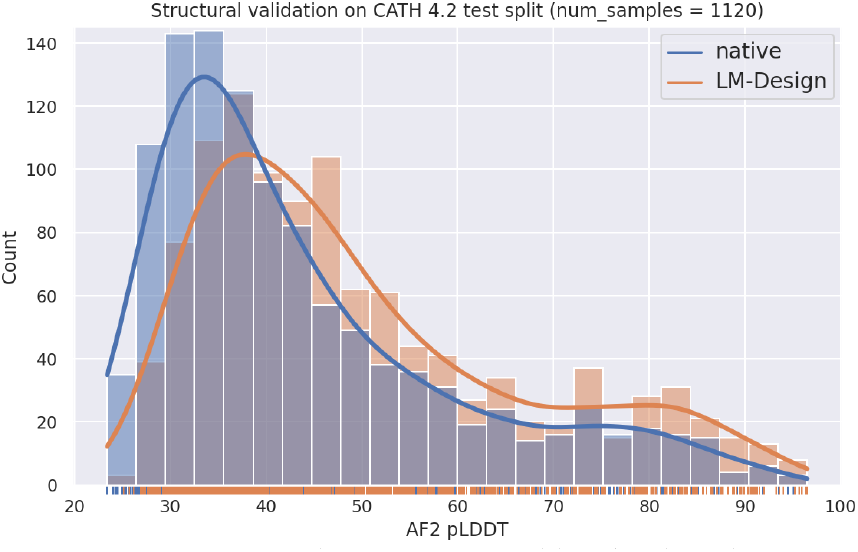
Structural validation measured by the distribution of AF2 pLDDT (→) for both native and LM-Design (ProteinMPNN) redesigned sequences.

#### 4.2.2. Studies on Inference

##### Iterative refinement gives rise to accurate sequence design

Since LM-Design is trained to denoise, we can exploit iterative decoding to progressively refine its predictions towards a better one (§3.2). As shown in **Fig. 3**, we find that even without iterative refinement LM-Design performs sufficiently well, while recycling the *p*LM-based decoder for LM-Design yields 1 ∼ 2% gains. This shows that iterative refinement is an effective strategy for sequence design if models are set up under a denoising learning scheme. Significant further improvement eliminates if iterating beyond 6 rounds, resulting in acceptable sampling efficiency.

**Figure 3.**
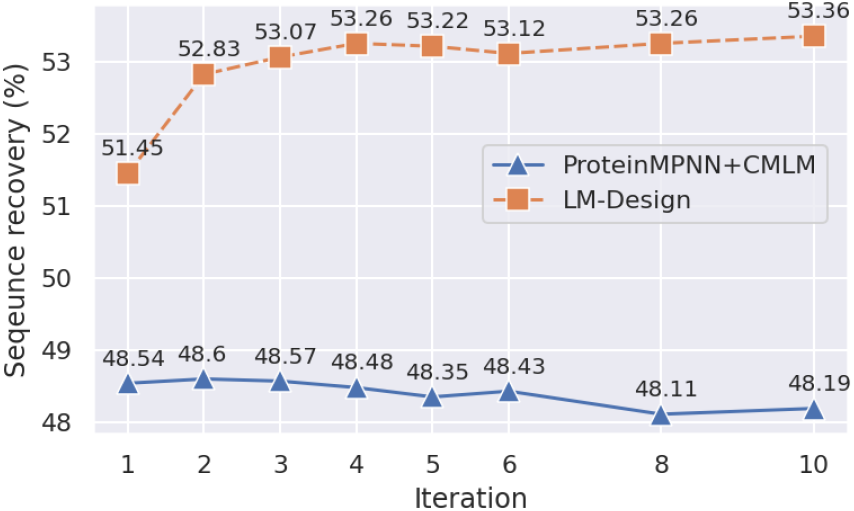
Effects of iterative refinement: sequence recovery *w*.*r*.*t*. decoding iterations.

##### Accuracy *vs*. diversity

While recent protein sequence design approaches have focused on maximizing native sequence recovery (and we also use it as our primary metric in experiments above), this is not necessarily optimal for actual protein design applications, for which novelty also matters (Dauparas et al., 2022). To this end, we manipulated the temperatures (*τ* ∈ [0.1, 0.5, 1.0, 1.2, 1.5]) to control the diversity of sampled sequences that are dissimilar to the native ones at different levels (as in §3.2), and evaluate design accuracy (in AF2 pLDDT) as a function of diversity. **Fig. 4** shows that **LM-D****esign** ***yields diverse yet more accurate designs*** over ProteinMPNN, manifesting the potential of practical values of LM-Design in real-world scenarios.

**Figure 4.**
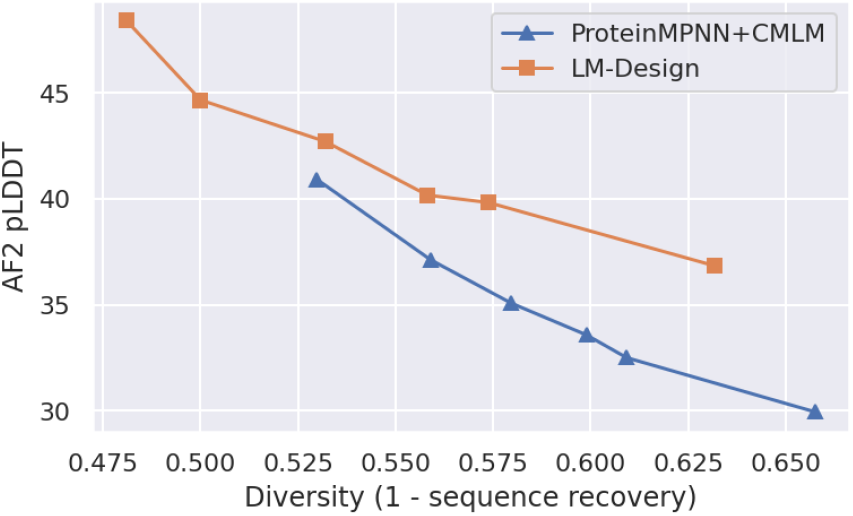
Pareto optimal points that represent the trade-off between accuracy (in AF2 pLDDT) and diversity, showing accurate and diverse predictions LM-Design can make.

#### 4.2.3. Does Scaling Help? Ablations on Scales of Structure Data and *p*LMs

##### LM-Design works well with data augmentation (Hsu et al., 2022) via incorporating predicted structures from AlphaFold 2

We perform different scales of data augmentation, the details of data processing are described in Appendix E. As the results in **Fig. 5** show that both of the methods obtain better results with 20k data augmentation. While ProteinMPNN+CMLM drops at 50k, LM-Design keeps increasing and finally drops at 100k. That is because LMDesign has 6.9M parameters while ProteinMPNN+CMLM only has 1.6M parameters. The same conclusion has also been mentioned in ESM-IF (Hsu et al., 2022), where models of less capacity cannot benefit from the increment of data.

**Figure 5.**
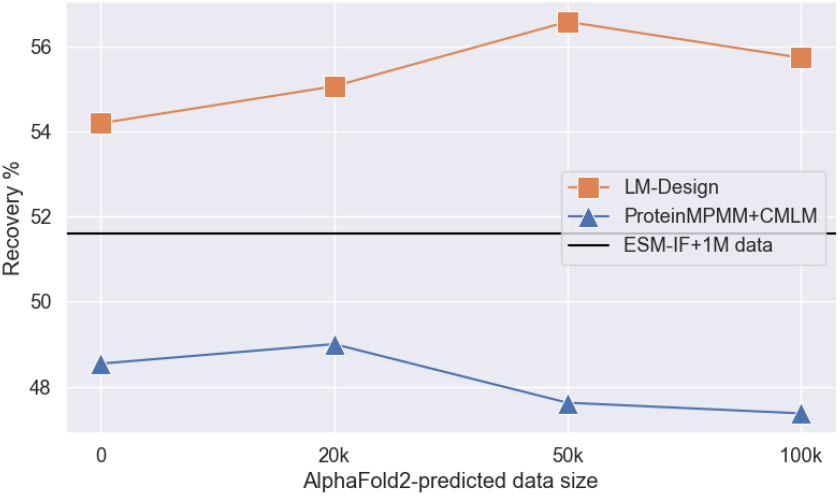
Performance *w*.*r*.*t. scales of augmented data*. Predicted data are produced by AlphaFold 2 from SWISS-PROT dataset.

##### LM-Design is scalable yet parameter-efficient: scaling law *w*.*r*.*t*. model sizes of *p*LMs using ESM-2 series (Lin et al., 2022)

To study the impact of the scale of *p*LMs, we switch LM-Design’s decoder from ESM-1b (145M) to ESM-2 series, with parameters (params) ranging from 8M to 3B. As shown in **Fig. 6**, the performance of LM-Design increase as model scaling, namely the larger the better, with a clear (log) scaling law exhibits. Such a scaling law agrees with large language models (LLMs) in general, which was initially observed and well-known in NLP (Kaplan et al., 2020). In addition, in contrast to existing strong systems (*e*.*g*., ESM-IF (Hsu et al., 2022) requires expensive training of all hundred millions of params in total), LM-Design is way more parameter-efficient only needing *<* 1% trainable params *w*.*r*.*t*. total params of the corresponding *p*LMs. In the extreme case, the largest ESM-2 3B-based variant has **22M** trainable out of **3B** total params (**0.07%**) and achieves the highest accuracy of 56.8%. We suggest that this strong connection between protein and natural languages on LLMs gives rise to exciting potentials to empower protein research with cutting-edge advances in general AI.

**Figure 6.**
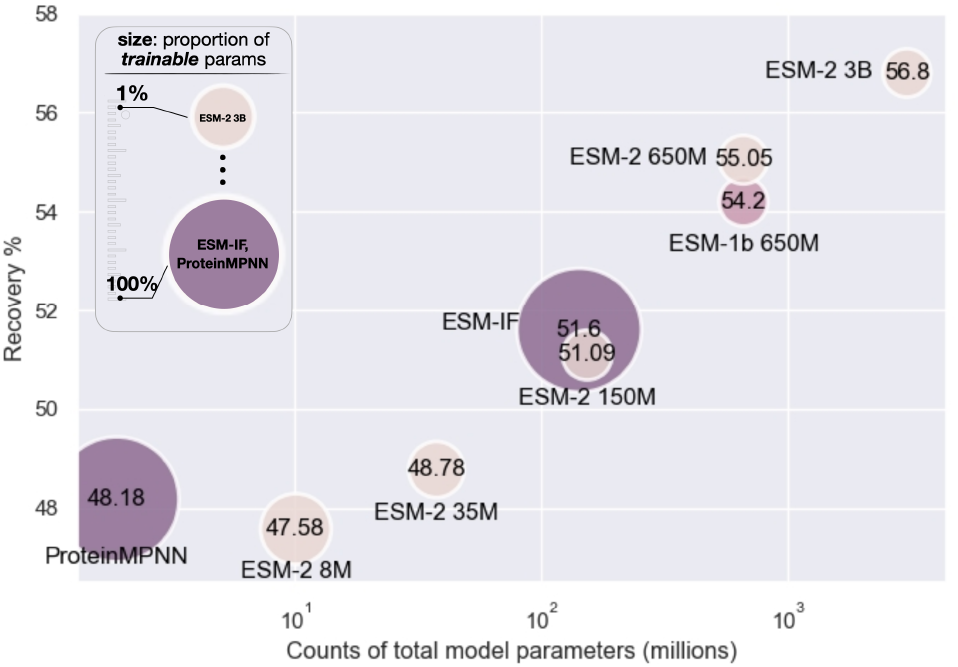
Performance *w*.*r*.*t. model scales of p*LM*s using ESM-2 series on CATH 4*.*3*. **X-axis**: counts of total model parameters; **Y-axis**: sequence recovery (↑); and the **bubble size**: proportions of trainable parameters over total model parameters (↓).

#### 4.2.4. How LM-Design Improves Protein Design? Studies on Protein Structures

##### LM-Design effectively exploits the potential of both structural and sequential capabilities

To further understand the action mechanism of LM-Design, we dissect its performance based on distinct structural contexts either with high structural constraints or low constraints. As shown in **Fig. 7A**, for single-chain proteins in the CATH dataset, structured-based ProteinMPNN shows high sequence recovery rates on structurally constrained residues in the folding core, while low recovery rates on structurally less-constrained residues on surface area and loops. Interestingly, LM-Design can effectively enhance the sequence recovery rates on structurally-constrained and less-constrained residues. Similar observations can be found for multichain complex proteins as shown in **Fig. 7B**. Although ProteinMPNN achieves high sequence recovery rates on folding core residues, it shows compromised performance on residues in the binding interface and exposed regions. LM-Design can generally improve sequence recovery rates in different structural contexts.

**Figure 7.**
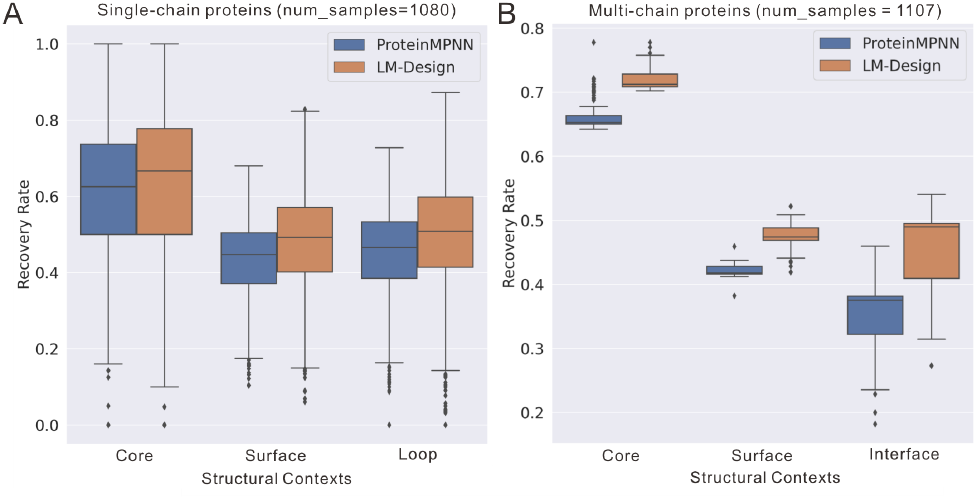
Comparison of sequence recovery *w*.*r*.*t. structural contexts regarding solvent accessible surface area and interaction interface*, on (A) CATH 4.2 single-chain proteins and (B) multichain protein complexes.

##### LM-Design is structurally sensitive

It is interesting to know whether the sequence-based protein language model LM-Design is sensitive to structure inputs. To evaluate this problem, we followed a previous biological study (Zhu et al., 2011), collected four proteins sharing similar structures but having distinct sequences with specific functions. We tested LM-Design’s performance using this case for designing specific functional sequences. As shown in **Fig. 8**, LMDesign can nicely predict functional specific sequences for each of the proteins, suggesting LM-Design is highly sensitive to structure variations.

**Figure 8.**
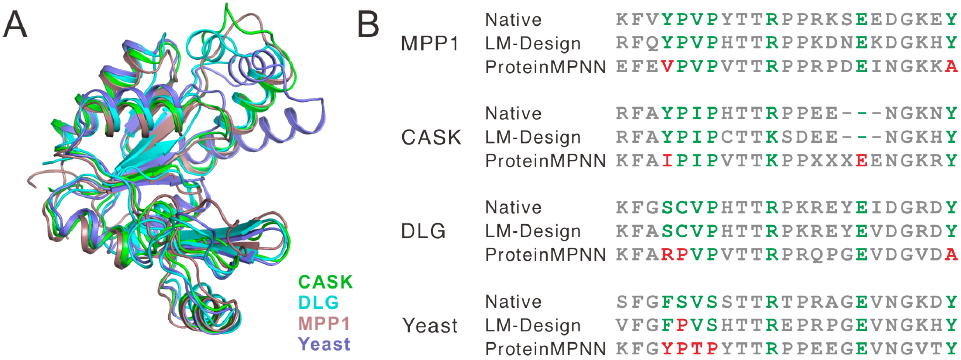
Case study to evaluate the structure sensitivity of the methods. (A) Structure alignments show highly similar structures of the four proteins. (B) Multiple sequence alignment. These distinct residues mediating specific biological functions are highlighted in **green**. The sequence predicted by LM-Design and ProteinMPNN were aligned to native sequences. The incorrect predictions are highlighted in **red**.

### 4.3. Zero-shot Generalization to Other Proteins

#### TS50 and TS500 datasets

TS50 and TS500 are commonly used independent test set to assess model generalization for proteins introduced by Li et al. (2014). We test LM-Design trained on CATH 4.2 and 4.3 respectively. As shown in **Tab. 3**, we can first find that models trained on CATH 4.3 seem to generalize better to these two datasets, and LMDesign consistently achieves the best. As for TS500, in particular, we observe that LM-Design improves PiFold (58.82% → 67.78%) and outperforms previous methods by a large margin. Notably, non-*p*LM-enpowered models (*e*.*g*., Pifold, though we re-implemented with comparable performance), sometimes assign way too higher probabilities to the incorrect amino acids, especially for the bad cases we observed in TS500, giving rise to enormous perplexity (marked as `n/a`). With the help of *p*LMs, LM-Design (Pi-Fold) goes back to a normal scale of perplexity, indicating the capabilities of *p*LMs again.

**Table 3.**
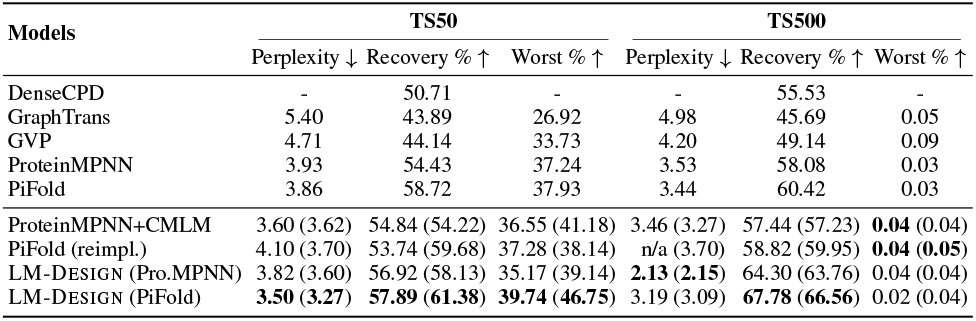
Performance comparison on the TS50 and TS500 datasets. We follow previous literature to mainly report the results using models trained on CATH 4.2. Results from models trained on CATH 4.3 are also provided (numbers in brackets). DenseCPD (Qi & Zhang, 2020) is a CNN-based (not GNN-based) approach.

#### *De novo* proteins

*De novo* protein design explores the full protein sequence space for a new range of folds with high stability. In the last decade, advanced computational methods have been developed to design protein structures with atomic-level accuracy (Huang et al., 2016). To test whether LMDesign can generalize to *de novo* proteins, we compiled a *de novo* protein dataset by collecting 134 *de novo* protein monomers with different folds from Protein Data Bank. The performance of LMDesign and ProteinMPNN was evaluated using this dataset showing an average recovery rate of 48.7%. LM-Design can recover the sequence with a significantly higher rate of 58.7%, suggesting a better generalization capability on designed proteins. Please refer to **Fig. 9** for detailed results.

**Figure 9.**
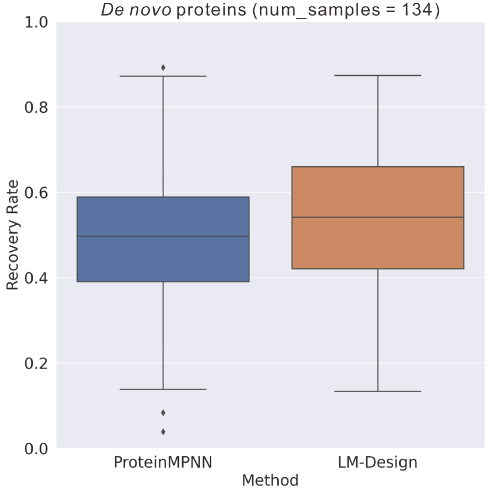
LM-Design shows a better generalization capability on *de novo* proteins than Protein-MPNN.

#### Antibody design

Designing targeted antibodies for different antigens is one of the potential treatments for many diseases that currently cannot be cured. Antibody design is formulated as sequence infilling for complementary-determining regions (CDRs) given contexts (*i*.*e*., antigen and antibody frameworks). However, the commonly-used metric of sequence recovery (*aka*. AAR, amino acid recovery) can be flawed as we often observed a “mode collapse” problem for previous approaches due to extremely limited antibody data. We hence design a package of five metrics regarding salient regions recovery, hallucinated sequence pattern, and entropy of predicted distribution for a more comprehensive evaluation (See Appendix §C.1 for details). We conducted two kinds of experiments on RAbD dataset (Adolf-Bryfogle et al. (2018)) in designing CDR-H3 sequences given either true complex structures, or predicted ones. (1) As for using true structures, we compare LM-Design with ProteinMPNN and MEAN (Kong et al., 2022), the SoTA neural antibody design approach using its `fixbb` mode. As shown in **Fig. 10A**, LM-Design outperforms antibody-specific MEAN model in terms of all evaluation aspects, showing that that models for general protein can effectively avoid mode failure while *p*LMs help facilitate antibody design. (2) In the case of predicted structures (**Fig. 10B**), the structures are predicted by MEAN. The performance of LM-Design and ProteinMPNN decreases significantly, implying they are fragile for predicted structures. In contrast, LM-Design + `eps`, where spatial perturbation is injected to the training structures (Dauparas et al., 2022), shows stronger robustness hence better performance, suggesting the need of counter-adversarial considerations for structure-based sequence models to enhance generalizability.

**Figure 10.**
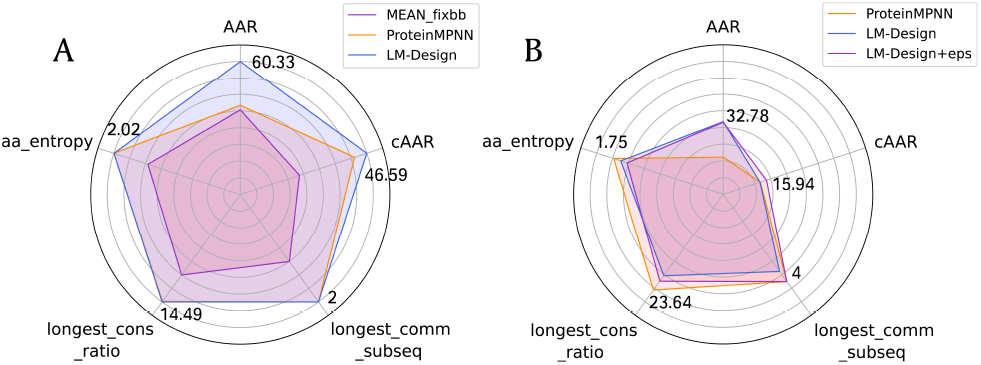
Performance comparison for antibody design. The farther away from the center of the circle, the better the performance. (A) Antibody sequence design based on true structure; (B) Antibody sequence design based on predicted structure

## 5. Discussions

In this paper, we demonstrate that language models are strong protein designers, and introduce LM-Design, a generic approach to enable *p*LMs to yield preferable protein sequences for given folds. We conduct a *structural surgery* on *p*LMs, where a lightweight structural adapter is implanted into *p*LMs and endows it with structural awareness. During inference, iterative refinement is performed to effectively optimize the generated protein sequences. Experiments show that LM-Design outperforms the state-of-the-art methods by a large margin. Extensive analyses verify that LM-Design can (1) effectively exploit both structural and sequential knowledge, (2) benefit from scaling data and size, and (3) generalize to other proteins (*e*.*g*., antibodies and *de novo* proteins).

Despite a leap of accuracy in terms of native sequence recovery in this study, there remain several limitations. The ultimate goal of computational protein design is to completely design *new* proteins based on a simple description of their target *function*. In machine learning, on the other hand, the dilemma of generalization (novelty) and memorization (recovery) is a long-standing question in evaluating generative models in general (van den Burg & Williams, 2021), namely, to what extent training data are memorized by the learning algorithm. As a result, the most crucial and immediate concern is whether a high native sequence recovery is a good indicator for successful (structure-based) protein design, which has also been noted and discussed in recent literature (Melnyk et al., 2022b).

For structure-based protein design, which lies in the current scope of this study, functions have already been determined by the given structures of interest^2^, the goal is therefore to explore protein sequence space to find novel sequences with optimal biochemical properties like stability and solubility. In this case, we have already showcased that, albeit *p*LMs being solely exposed to tens of millions *native* protein sequences, LM-Design is capable of leveraging the patterns learned by *p*LMs and combining them to yield accurate yet diverse (**Fig. 4**) and structurally valid (**Fig. 2**) protein sequences for given folds.

As for protein design at large, novel structures distant from natural proteins are often more emphasized (Watson et al., 2022). Recently, Lin et al. (2022) show that sequential evolutionary knowledge learned by *p*LMs corresponds deeply to protein structures and thus materializes protein structure prediction from single sequences. In a concurrent work, Verkuil et al. (2022) further demonstrate that *p*LMs at scale can synthesize *de novo* proteins on the basis of the deep grammars learned from large-scale native sequences, generalizing beyond natural proteins, at a high experimental success rate. Likewise, in the recent advances in generative AI algorithms in general, *e*.*g*., VAE (Kingma & Welling, 2014), autoregressive models (Bahdanau et al., 2015; Vaswani et al., 2017), and diffusion probabilistic models (Sohl-Dickstein et al., 2015; Ho et al., 2020; Rombach et al., 2022), maximum likelihood estimation or its surrogates is the primary protocol for learning from massive native data by reconstruction. Once they have learned to sufficiently recover native data, they can synthesize completely new data, yielding impressive results and thus driving the recent surge of generative AI, both in academia and industry, and with great potential even in the field of protein (structure) design (Watson et al., 2022).

All of these show that, in our opinion, a good recovery is a prerequisite for novelty: a generative model with good adherence to the native data in its vastness and the ability to recover them (and their in-domain held-out set), if over-fitting does not occur, captures the underlying data distribution in a great extent, hence has the capability to generalize and synthesize *de novo* data from the patterns and “grammars” they learned.

Based on the above discussions, we suggest that the good performance of LM-Design can be attributed to the aforementioned reasons. We expect LM-Design can further benefit from more ingenious decoding/sampling algorithms for better ***designability*** (*e*.*g*., being more diverse, constrained, controllable, and programmable), integrating with structure prediction models (Jumper et al., 2021; Baek et al., 2021) for the purpose of end-to-end protein ***structure-sequence codesign*** (Jin et al., 2021), incorporating stochastic dynamics and variational Bayes to ***better capture multimodal distributions*** as in VAEs/diffusion models, and even utilizing protein analogs of ***in-context learning*** (Wei et al., 2022a) or ***chain-of-thoughts (CoT) prompting*** (Wei et al., 2022b; Fu et al., 2022) in LLMs for natural languages. Moreover, as the scaling laws link performance with the computing, data, and model size in LLMs in general and proteins, we can be optimistic about the further breakthroughs in generative protein research enabled by the advances of scaled general sequence learning algorithms and their empowered protein applications, together with the progress in the fundamental protein geometry and structure learning. We leave these exciting directions as future work.

## Acknowledgements

We would like to especially thank Dr. Hang Li for insightful discussions on the project and feedback on the manuscript that help shape this study. We thank Siyu Long, Xiaoqin Tan, and Yu Bao for their valuable comments. We also thank Chloe Hsu for addressing our questions about the experimental evaluation in Hsu et al. (2022).

## A. Model Architecture of LM-DESIGN

The overall architecture of LM-Design is constituted by three components, *i*.*e*., a structure encoder, a *p*LM as sequence decoder, and a structural adapter that bridges both. We directly utilized the established structure models (architectures, hyper-parameters, *etc*.) from literature to parameterize the structure encoder, in which we showcased ProteinMPNN (Dauparas et al., 2022) as our primary running example, and also investigated PiFold (Gao et al., 2022) and GVP-TransformerEncoder (Hsu et al., 2022) (we discarded the TranformerDecoder in GVP-Transformer and only kept the encoder part as our only purpose is encoding protein structure). For *p*LMs as the sequence decoder side, we primarily used the ESM-1b 650M (Rives et al., 2019), as well as ESM-2 series (Lin et al., 2022), with their pretrained model weights.

Note that inventing better protein structure encoding modules or sequence pretraining/representation learning approaches is not the focus thus beyond the scope of our study. We built LM-Design on top of these popular and publicly available models as backbones. We strongly encourage readers to check the details of these models in their original papers. In addition, we believe LM-Design as a generic framework can further be improved by any future advances in the backbones in terms of both protein structure and sequence learning.

Here we elaborate on the design of the structural adapter. **Fig. 11** shows one instantiation we used in this paper, despite there possibly existing many others to explore, the structural adapter composes a multihead attention (Multihead Attn) that queries structure information from the structure encoder, followed by a *bottleneck* feedforward network (FFN) to impose non-linearity and abstract features/representations. The bottleneck architecture of the FFN adheres to the best practice from Houlsby et al. (2019) and many follow-ups of it. Most hidden dimensions of Multihead Attn and FFN are determined by the instance of structure encoder and *p*LM we used, while the intermediate dimension of the bottleneck FFN was set to be half of the model dimension. RoPe (Su et al., 2021) was used the supplement Multihead Attn for better modeling of positional information. In all our experiments, only one structural adapter was placed after the last layer of *p*LM, although layer-wise placement is also an obvious alternative and we leave it for future study.

**Figure 11.**
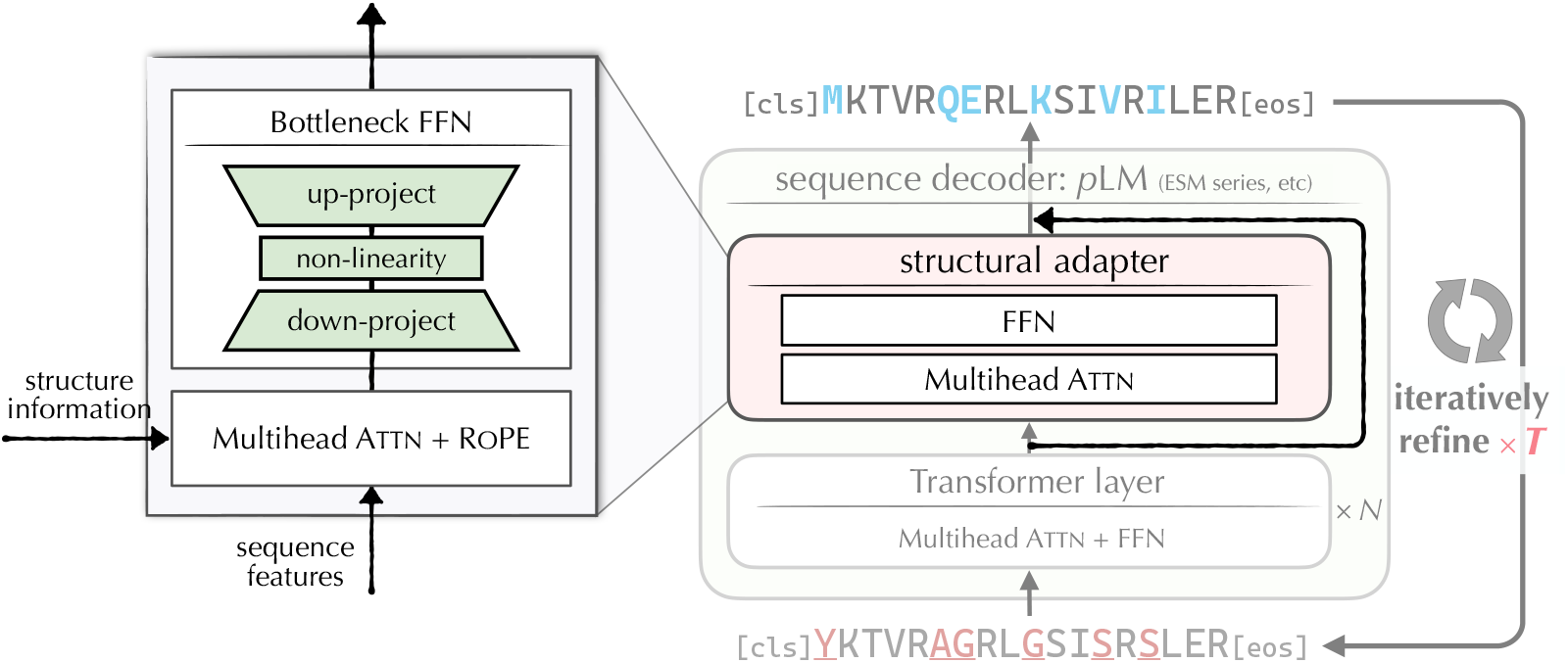
Illustration of our instantiation of the structural adapter

## B. Additional Details on Benchmarking Evaluations

### B.1. Single-chain Protein Design on CATH Datasets

#### Datasets

We mainly compared LM-Design against recent strong baselines on the CATH 4.2 (Orengo et al., 1997) dataset, using the same data splits as the compared systems^3^, *e*.*g*., Structured Transformer (Ingraham et al., 2019), GVP (Jing et al., 2020), ProteinMPNN (Dauparas et al., 2022), and PiFold (Gao et al., 2022), where proteins were partitioned by the CATH 4.2 topology classification, resulting in 18024 proteins for training, 608 proteins for validation, and 1120 proteins for testing. To compare with ESM-IF (Hsu et al., 2022), we also conducted evaluations on CATH 4.3^4^, wherein 16153 structures are assigned to the training set, 1457 to the validation set, and 1797 to the test set.

#### Implementation, training, and metrics

The models were trained up to 100 epochs by default using the Adam optimizer on NVIDIA V100s. We used the same training settings as ProteinMPNN (Dauparas et al., 2022), where the batch size was set to approximately 6000 residues, and Adam optimizer (Kingma & Ba, 2015) with `noam` learning rate scheduler (Vaswani et al., 2017) was used. We follow the previous studies to report perplexity and *median* recovery scores on short proteins (length ≤ 100), single-chain proteins (labeled with 1 chain in CATH, different from the single-chain concept which is in contrast with protein complex), and all proteins settings.

### B.2. Multi-chain Protein Complex Design

#### Dataset and setting

We used the multi-chain complex dataset curated by Dauparas et al. (2022), which collected protein assemblies in the PDB (as of Aug 02, 2021) determined by X-ray crystallography or cryoEM to better than 3.5Å resolution and with less than 10,000 residues^5^. Sequences were clustered at 30% sequence identity cutoff using `mmseqs2` (Steinegger & Söding, 2017) resulting in 25,361 clusters. Those clusters were split randomly into three groups for training (23,358), validation (1,464), and testing (1,539), ensuring that none of the chains from the target chain or chains from the biounits of the target chain would be in the other two groups. We kept the training setting the same as that of the single-chain scenario.

## C. Antibody Design

### C.1. Task Definition, Datasets, and Evaluation Metrics

#### Task definition

We design two types of experiments.

- **Sequence design on true structures**. In this experiment, we set the antibody sequence design task as a fixed-backbone protein design task in accordance with the training task of LM-Design, wherein each model is asked to design the CDR-H3 region sequence of the antibody-based on the true antigen-antibody complex structure^6^.
- **Sequence design on predicted structures**. In this experiment, we examine the performance of the model when facing a more real situation, lacking the real structure of the antibody. All the models are asked to predict the CDR-H3 sequence based on the predicted CDR structure. We use MEAN (Kong et al., 2022), the SoTA method in the designing of antibody generate the structure of the CDR region. Specifically, we modify the MEAN by initializing the CDR sequence using AbLang (an antibody pre-training model, Olsen et al. (2022)) to obtain a fine predicted CDR-H3 structure (RMSD 1.81 → 1.70).

It should be noted that although antibodies are also proteins, antibody design, i.e. CDR design, is very different from protein design. Protein sequence diversity is evolutionary in origin, whereas CDR sequence diversity is derived from V(D)J recombination and somatic hypermutation in B cells, which indicates that CDR sequences are random to some extent.

#### Datasets

We follow the dataset adopted by most antibody design work. 60 antigen-antibody complexes from RAbD dataset were used to test LM-Design’s design capability on the antibody sequence when given the structure of the antigen-antibody complex, and all antibodies are renumbered under the IMGT scheme (Lefranc et al. (2003)).

#### Evaluation metrics

AAR is the most important metric in antibody design. However, because of the conservation of amino acids at both ends of the CDR region and the insensitivity of sequence recovery to short sequences, AAR may not reflect the true performance of the model. For example, similar antibodies like “`ARD G* Y* FDY`” are generated for different antigens, it fails to targeted design but still achieves good accuracy (37.5% AAR on RAbD dataset).

As a result, to evaluate the model’s performance more comprehensively, we introduced four additional metrics to evaluate the generated CDR-H3 sequence in addition to widely used AAR. Concretely, `longest comm subseq` is the length of the longest common subsequence in at least 30% of generated sequences, indicating the lack of diversity in the generated sequence, lower is better; `longest cons ratio` is the proportion of the longest consecutive repeat sequence in the generated CDR sequence, indicating the occurrence of polyY and polyG in the generated sequence, lower is better. `aa entropy` is the average entropy of each amino acid type in each generated CDR sequence, indicating the diversity of amino acid types in the CDR sequence, higher is better. `cAAR` is the `AAR` calculated at the actual interaction position, which can better reflect the model’s understanding of complex protein structures and interactions, higher is better.

### C.2. Additional Results

#### Sequence design on true structures

MEAN is the method specially designed for co-designing CDR structures and sequences, and we used open-source code of MEAN and data (data were filtered down from 3127 to 2901 to remove the proteins that lacked atomic information) to retrain `fixbb` mode MEAN in this experiment, the training config is consistent with the original MEAN. The experiment results are shown in **Tab. 4**. In the first experiment, LM-Design and ProteinMPNN+CMLM achieve far better performance than the original ProteinMPNN, and also outperformed MEAN in `fixbb` mode. Among all the five metrics, the performance gap between LM-Design and ProteinMPNN+CMLM in `cAAR` is greater than the others. We believe that the language model in LM-Design may “forget” certain structural information, resulting in a significant decline in `cAAR`, which is closely related to structure. To avoid this “forgetting” of the structure, we add the output logits of the structure encoder to LM-Design’s output logits to enhance the impact of the structure on the predicted sequence. Finally, LM-Design +`Enc_Logits` achieves the best performance in four of the five metrics.

**Table 4.**
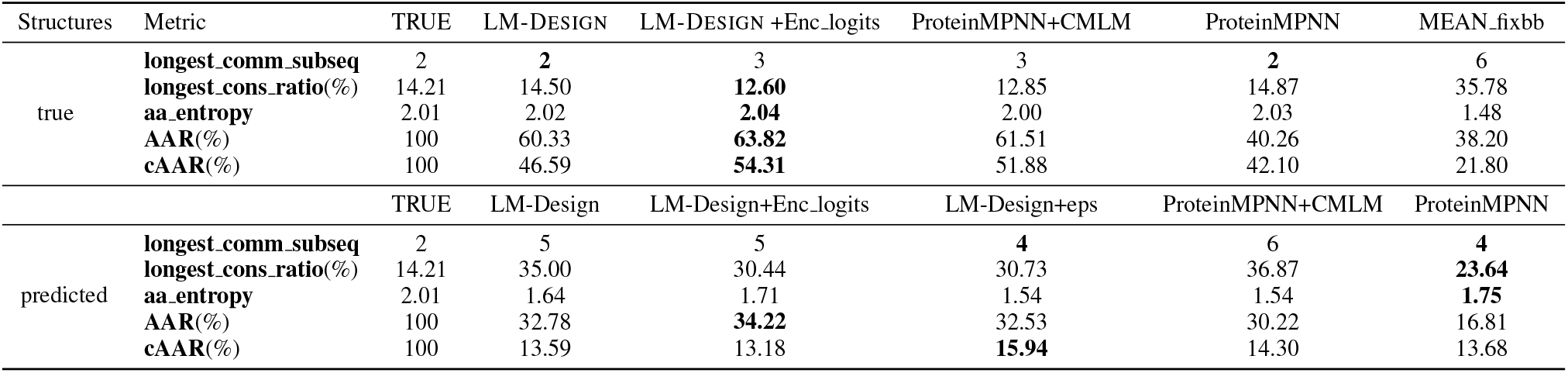
Performance on RAbD dataset. The scores of the original ProteinMPNN were obtained using its publicly available model checkpoint.

| Structures | Metric | TRUE | LM-DESIGN | LM-DESIGN +Enc_logits | ProteinMPNN+CMLM | ProteinMPNN | MEAN_fixbb |
| --- | --- | --- | --- | --- | --- | --- | --- |
| true | **longest_comm_subseq** | 2 | **2** | 3 | 3 | **2** | 6 |
|  | **longest_cons_ratio(%)** | 14.21 | 14.50 | **12.60** | 12.85 | 14.87 | 35.78 |
|  | **aa_entropy** | 2.01 | 2.02 | **2.04** | 2.00 | 2.03 | 1.48 |
|  | **AAR(%)** | 100 | 60.33 | **63.82** | 61.51 | 40.26 | 38.20 |
|  | **cAAR(%)** | 100 | 46.59 | **54.31** | 51.88 | 42.10 | 21.80 |
|  |  | TRUE | LM-Design | LM-Design+Enc_logits | LM-Design+eps | ProteinMPNN+CMLM | ProteinMPNN |
| predicted | **longest_comm_subseq** | 2 | 5 | 5 | **4** | 6 | **4** |
|  | **longest_cons_ratio(%)** | 14.21 | 35.00 | 30.44 | 30.73 | 36.87 | **23.64** |
|  | **aa_entropy** | 2.01 | 1.64 | 1.71 | 1.54 | 1.54 | **1.75** |
|  | **AAR(%)** | 100 | 32.78 | **34.22** | 32.53 | 30.22 | 16.81 |
|  | **cAAR(%)** | 100 | 13.59 | 13.18 | **15.94** | 14.30 | 13.68 |

#### Sequence design on predicted structures

In this experiment, The original ProteinMPNN achieves the best performance in three metrics of diversity, but shows great disadvantages in AAR (**Tab. 4**). LM-Design and LM-Design +Enc logits perform higher AAR, while ProteinMPNN+CMLM gets better cAAR. However, all methods show a huge performance degradation when using the predicted structure instead of the true structure, and this phenomenon shows the sensitivity of antibody design tasks to structural changes. We conducted preliminary experiments and try to mitigate this degradation, as we usually do not know the true structure of the CDR region in actual antibody design tasks. Specifically, we inject spatial perturbation into LM-Design’s training structure and hope that this will help the model to be more robust when structures have uncertainty. The resulting model (LM-Design +eps) achieves significant improvements in cAAR, proving that tolerance for structural bias is helpful for the actual antibody sequence design task.

In general, in antibody sequence design tasks, LM-Design and ProteinMPNN+CMLM can greatly improve the accuracy of the generated sequence at the expense of limited diversity. More specifically, LM-Design discards some perception of structure awareness based on ProteinMPNN+CMLM and gains a stronger ability to model sequences.

## D. *De novo* Protein Design

### Task definition

The aim of computational protein design is to invent novel protein molecules with desired structures and functions (Huang et al., 2016). In this experiment, we evaluate the performance of the model to generalize to fixed-backbone sequence design on *de novo* proteins.

### Dataset and evaluation metrics

To test LM-Design’s generalization capability, we compiled 134 single-chain *de novo* proteins (length ≤ 30) from the PDB data bank. The structures of these samples were determined by X-ray crystallography or cryo-EM to better than 3.5Å resolution The sequences were clustered at 30% sequence identity cutoff using mmseqs2 (Steinegger & Söding, 2017). We follow the other design tasks using median recovery scores for the test.

## E. Data Augmented Training with Predicted Structures from AlphaFold 2

### Task definition

Due to limited experimentally determined protein structure data and the surge of protein structure prediction models (*e*.*g*., AlphaFold 2), a natural idea for better protein inverse folding is to use the protein structure data predicted by AlphaFold 2 for data augmentation, which is similar to the back-translation in NLP area. For the protein design task, 𝒳 and 𝒴 are the sets of protein structure coordinates and protein sequences. The goal is to learn a mapping *f* : 𝒳 → 𝒴 while AlphaFold 2 has learned a mapping *g* : 𝒴 → 𝒳. We denote those protein sequences without structures as 𝒰 _𝒴_⊂ 𝒴. For any sequence *y*_*u*_ ∈ 𝒰_𝒴_, we can predict its structure 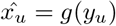 and add 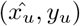 to our training set.

### Datasets

UniRef50 (Suzek et al., 2015) is a sequence database that has over 50 million clusters at 50% sequence identity, ESM-IF (Hsu et al., 2022) predicts structures of 12 million sequences in UniRef50 using AlphaFold 2. Since our main goal is to prove the feasibility of combining language model design and data augmentation, we use SWISS-PROT (Boeckmann et al., 2003) in our experiment. SWISS-PROT is a curated protein sequence database that strives to provide a high level of annotation, a minimal level of redundancy, and a high level of integration with other databases.

In order to prevent data leakage introduced by data augmentation, we need to exclude proteins that have the same fold as the proteins in validation and test splits. We annotated SWISS-PROT sequences with CATH fold classification according to the Gene3D (Lees et al., 2011) database with `hmmsearch` (Potter et al., 2018). We then perform filtering based on all CATH folds from validation and test splits and randomly select a sub-set of 100, 000 examples from the filtered dataset. We fix the random seed and randomly select 20, 000 and 50, 000 sequences for different scales of data augmentation.

## F. Structural Contexts Analysis

### F.1. Methods

In order to understand how LM-Design improves protein design, we analyze the performance based on dissected structural contexts (folding core residues, surface exposed residues, residues located on loops, and residues located on complex interfaces). The dictionary of structural contexts is obtained by using the widely used DSSP (Kabsch & Sander, 1983) and `biopython` (Cock et al., 2009) tools. Solvent accessible surface area analysis from DSSP calculates the ratio of solvent accessible surface area to the maximum possible solvent accessible surface area for each residue. A cutting threshold of 0.1 is chosen to classify residues located in the folding core and on the exposed surface. DSSP is also used to provide secondary structure labels. Those residues showing turn, bend, and none secondary structure patterns are labeled as loops in this study. For interaction interface analysis, we utilize InterfaceBuilder from Biopython. When the structural context label is obtained for each residue, the average recovery rate is calculated on PDB structures.

### F.2. Performance Dissection Results

#### Protein design

As the results shown in **Fig. 7**. As expected, residues in the folding core with more structural constraints display better sequence recovery rates than surface exposed and loop residues using both the structure-based ProteinMPNN and sequence-based LM-Design. Notably, the residues in the binding interface of multichain proteins show a poor recovery rate, suggesting the limited representation capability of existing structure-based methods in interaction interface design. LM-Design can generally improve sequence recovery rates in different structural contexts, including the interaction interfaces.

#### *p*LMs model scales

To study the impact of the model scale of *p*LMs on single-chain and multi-chain protein design, we dissect their performance in different structural contexts. As shown in **Fig. 12**A, the recovery rate of residues in the folding core increased from 65.2% (ESM-1b 650M) to 67.0% (ESM-2 650M), and finally to 69.3% using ESM-2 3B. For multi-chain proteins, ESM-2 3B also achieves the best recovery rate among different structure contexts (**Fig. 12**B). Therefore, the performance improvement can be found as a general trend for different structural contexts in sequence recovery when using *p*LMs with a larger scale of parameters. This result suggests larger *p*LMs models benefit from the training on the larger-scale protein sequence.

**Figure 12.**
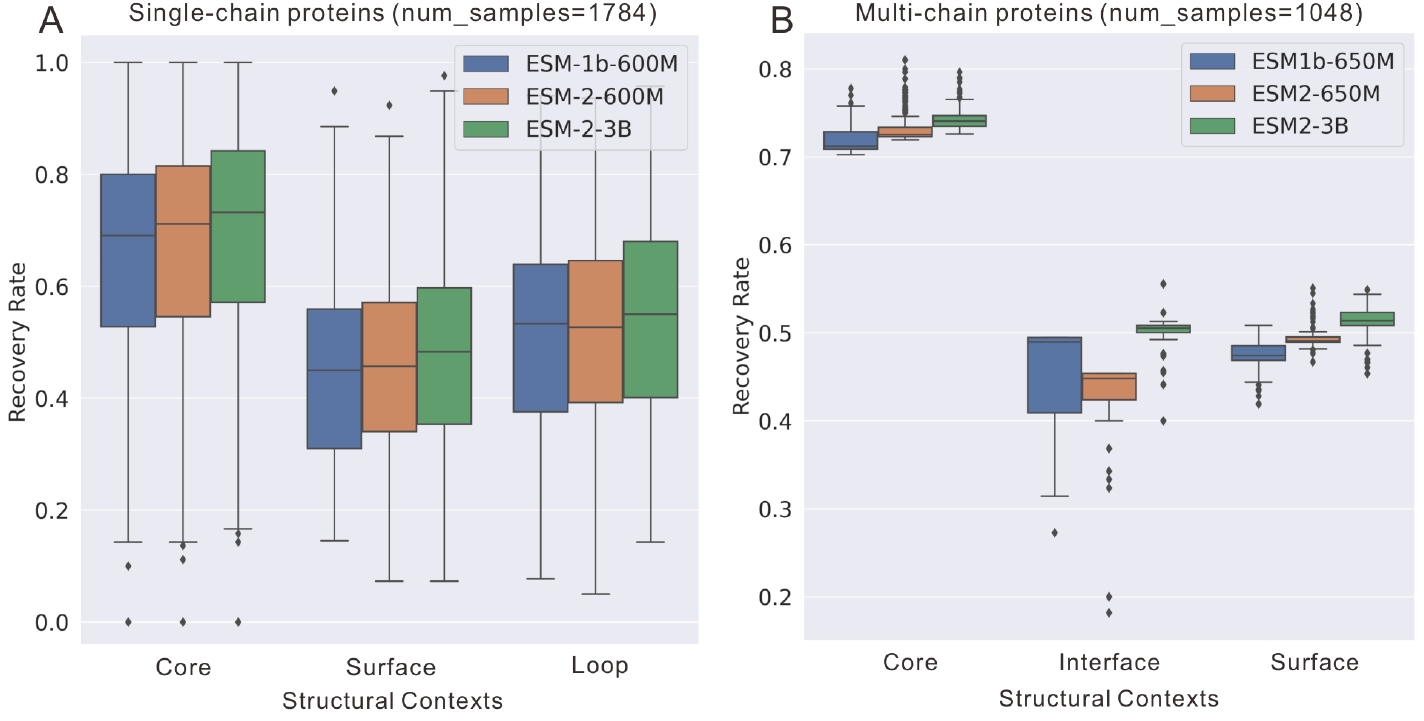
Comparison of sequence recovery by structural contexts for evaluation of the model scale of *p*LMs. (A) Single-chain results. (B) Multiple-chain protein results.

## G. Related Work

### Structure-based Protein Sequence Design with Deep Generative Models

Deep generative modeling typically formulates structure-based protein sequence design as a conditional sequence generation problem, wherein protein 3D structures can typically be represented as a *k*-NN graph (Ingraham et al., 2019). Several graph neural networks (GNNs) can be applied in this case to derive protein structural features. The protein graph establishes edge features between adjacent residues and encodes residue information as node features. The graph attention encoder and autoregressive decoder are used by GraphTrans (Ingraham et al., 2019) for protein design. The novel geometric vector perceptrons which take into account both scalar and vector features, GVP (Jing et al., 2020) improve further performance. Global graph attention for protein design is further introduced by GCA (Tan et al., 2022) to capture long-range information. Recently, ProteinMPNN (Dauparas et al., 2022) and PiFold (Gao et al., 2022) introduce more complicated protein features and expressive GNNs and gain significant improvements. Besides the primary generative purpose, Yang et al. (2022b) attempt to use this task as a proxy for protein (structure-aware) representation learning. In this study, we develop LM-Design on top of the powerful structural capability of ProteinMPNN and PiFold, while LM-Design can further benefit from future progress in deep geometric learning for proteins.

There, however, remains a crucial and severe issue that limits deep generative models for structure-based protein design is the lack of sufficient protein structure data. To address this, ESM-IF (Hsu et al., 2022) achieves dramatic improvements with effective data augmentation. Hsu et al. (2022) augments 16k experimental structures from CATH with 12 millions additional predicted structures “back-translated” (Sennrich et al., 2016) by AlphaFold 2 (Jumper et al., 2021). By further scaling the model to over a hundred million parameters, the ESM-IF model (Hsu et al., 2022) exhibits the state-of-the-art performance by a gain of over 10% sequence recovery (*i*.*e*., 51.6% on CATH 4.3), compared to the scenarios where only experimental data is considered. Note that LM-Design is orthogonal to such an effective data augmentation, which has been empirically verified by our experiment. By combining our modeling advances with data augmentation, we can take the best of both worlds to push the boundary of structure-based protein design.

### Protein Language Models

There is growing interest in developing protein language models (*p*LMs) at the scale of evolution due to the abundance of 1D amino acid sequences, such as the series of ESM (Rives et al., 2019; Lin et al., 2022), TAPE (Rao et al., 2019), ProtTrans (Elnaggar et al., 2021), PRoBERTa (Nambiar et al., 2020), PMLM (He et al., 2021), ProteinLM (Xiao et al., 2021), PLUS (Min et al., 2021), Adversarial MLM (McDermott et al., 2021), ProteinBERT (Brandes et al., 2022), CARP (Yang et al., 2022a) in masked language modeling (MLM) fashion, ProtGPT2 (Ferruz et al., 2022) in causal language modeling fashion, and several others (Melnyk et al., 2022a; Madani et al., 2021; Unsal et al., 2022; Nourani et al., 2021; Lu et al., 2020; Sturmfels et al., 2020; Strodthoff et al., 2020). These protein language models are able to generalize across a wide range of downstream applications and can capture evolutionary information about secondary and tertiary structures from sequences alone. They have recently been demonstrated with strong capabilities in uncovering protein structures (Lin et al., 2022), predicting the effect of sequence variation on function (Meier et al., 2021), antibody infilling (Melnyk et al., 2022a) and many other general purposes (Rives et al., 2019).

Concurrently, Verkuil et al. (2022) demonstrate that *p*LMs at scale can generalize beyond natural proteins to generate *de novo* proteins, and validate their hypothesis *in silico* and experimentally in great detail, in which *p*LMs are capable of even designing protein structure even though they only trained on sequences.

### Parameter-efficient Tuning for Language Models in NLP

The ability to effectively tune and adapt LLMs with few trainable parameters is crucial for modern NLP research and applications, because it is becoming increasingly impractical to train and store full copies of large LLMs for a variety of downstream tasks. Based on whether new trainable parameters are added, existing methods for LLM adaptation can be roughly divided into two categories. For example, one could train a fraction of the model’s parameters, such as the prediction head and bias terms (Lee et al., 2019) or introduce task-specific parameters to various LLM components (Li & Liang, 2021). Among them, Houlsby et al. (2019) have studied the choice of adapter architectures and the placement of them within LLMs in great detail. They discover that a stack of bottleneck networks, which only adds a few extra parameters to the network, performs well. The majority of the subsequent research (Pfeiffer et al., 2020a;c) and community-contributed framework, the AdatperHub (Pfeiffer et al., 2020b), are inspired by this design.

However, there yet remains less effort in transferring protein large language models for downstream important protein problems of interest, especially for the generative setting such as protein design. To the best of our knowledge, LMDesign is among the first attempts to steer *p*LMs to perform protein design, surpassing all previous non-*p*LMs empowered approaches for this task, suggesting the call for more attention in the power and potential of *p*LMs for protein research.

## Footnotes

1 A “full-mask” initialization also works well but considerably sub-optimal.

2 In the setting of fixed backbone sequence design, we assume structure design has been done beforehand as a prerequisite.

3 https://github.com/jingraham/neurips19-graph-protein-design/blob/master/data/README.md

4 https://github.com/facebookresearch/esm/blob/main/examples/inverse_folding/README.md

5 https://github.com/dauparas/ProteinMPNN/blob/main/training/README.md

6 CDRs, short for complementary-determining regions, are a few special regions on the antibody, with high variability and true interaction with the antigen, which determines the specificity of the antibody. CDR-H3 is the most important CDR.

## References

Adolf-Bryfogle, J., Kalyuzhniy, O., Kubitz, M., Weitzner, B. D., Hu, X., Adachi, Y., Schief, W. R., and Dunbrack Jr, R. L. Rosettaantibodydesign (rabd): A general framework for computational antibody design. PLoS computational biology, 14(4):e1006112, 2018.

Alford, R. F., Leaver-Fay, A., Jeliazkov, J. R., O’Meara, M. J., DiMaio, F. P., Park, H., Shapovalov, M. V., Renfrew, P. D., Mulligan, V. K., Kappel, K., et al. The rosetta all-atom energy function for macromolecular modeling and design. Journal of chemical theory and computation, 13 (6):3031–3048, 2017.

Baek, M., DiMaio, F., Anishchenko, I., Dauparas, J., Ovchinnikov, S., Lee, G. R., Wang, J., Cong, Q., Kinch, L. N., Schaeffer, R. D., et al. Accurate prediction of pro-tein structures and interactions using a three-track neural network. Science, 373(6557):871–876, 2021.

Bahdanau, D., Cho, K., and Bengio, Y. Neural machine translation by jointly learning to align and translate. In Bengio, Y. and LeCun, Y. (eds.), 3rd International Confer-ence on Learning Representations, ICLR 2015, San Diego, CA, USA, May 7-9, 2015, Conference Track Proceedings, 2015. URL http://arxiv.org/abs/1409.0473.

Boeckmann, B., Bairoch, A., Apweiler, R., Blatter, M.-C., Estreicher, A., Gasteiger, E., Martin, M. J., Michoud, K., O’Donovan, C., Phan, I., et al. The swiss-prot pro-tein knowledgebase and its supplement trembl in 2003. Nucleic acids research, 31(1):365–370, 2003.

Brandes, N., Ofer, D., Peleg, Y., Rappoport, N., and Linial, M. Proteinbert: a universal deep-learning model of pro-tein sequence and function. Bioinformatics, 38(8):2102–2110, 2022.

Brown, T., Mann, B., Ryder, N., Subbiah, M., Kaplan, J. D., Dhariwal, P., Neelakantan, A., Shyam, P., Sastry, G., Askell, A., et al. Language models are few-shot learners. 2020.

Cock, P. J., Antao, T., Chang, J. T., Chapman, B. A., Cox, C. J., Dalke, A., Friedberg, I., Hamelryck, T., Kauff, F., Wilczynski, B., et al. Biopython: freely available python tools for computational molecular biology and bioinformatics. Bioinformatics, 25(11):1422–1423, 2009.

Dahiyat, B. I. and Mayo, S. L. Probing the role of packing specificity in protein design. Proceedings of the National Academy of Sciences, 94(19):10172–10177, 1997.

Dauparas, J., Anishchenko, I., Bennett, N., Bai, H., Ragotte, R. J., Milles, L. F., Wicky, B. I., Courbet, A., de Haas, R. J., Bethel, N., et al. Robust deep learning–based pro-tein sequence design using proteinmpnn. Science, 378 (6615):49–56, 2022.

Devlin, J., Chang, M.-W., Lee, K., and Toutanova, K. BERT: Pre-training of deep bidirectional transformers for lan-guage understanding. In Proceedings of the 2019 Con-ference of the North American Chapter of the Associa-tion for Computational Linguistics: Human Language Technologies, Volume 1 (Long and Short Papers), pp. 4171–4186, Minneapolis, Minnesota, June 2019. Asso-ciation for Computational Linguistics. doi: 10.18653/v1/N19-1423. URL https://www.aclweb.org/anthology/N19-1423.

Elnaggar, A., Heinzinger, M., Dallago, C., Rehawi, G., Wang, Y., Jones, L., Gibbs, T., Feher, T., Angerer, C., Steinegger, M., et al. Prottrans: Toward understanding the language of life through self-supervised learning. IEEE transactions on pattern analysis and machine intelligence, 44(10):7112–7127, 2021.

Ferruz, N. and Höcker, B. Controllable protein design with language models. Nature Machine Intelligence, 4(6): 521–532, 2022.

Ferruz, N., Schmidt, S., and Höcker, B. Protgpt2 is a deep unsupervised language model for protein design. Nature communications, 13(1):4348, 2022.

Fu, Y., Peng, H., Sabharwal, A., Clark, P., and Khot, T. Complexity-based prompting for multi-step reasoning. arXiv preprint arXiv:2210.00720, 2022.

Gao, Z., Tan, C., and Li, S. Z. Pifold: Toward effec-tive and efficient protein inverse folding. arXiv preprint arXiv:2209.12643, 2022.

Ghazvininejad, M., Levy, O., Liu, Y., and Zettlemoyer, L. Mask-predict: Parallel decoding of conditional masked language models. In Proceedings of the 2019 Confer-ence on Empirical Methods in Natural Language Pro-cessing and the 9th International Joint Conference on Natural Language Processing (EMNLP-IJCNLP), pp. 6112–6121, Hong Kong, China, November 2019. As-sociation for Computational Linguistics. doi: 10.18653/v1/D19-1633. URL https://www.aclweb.org/anthology/D19-1633.

Gu, J., Bradbury, J., Xiong, C., Li, V. O. K., and Socher, R. Non-autoregressive neural machine translation. In 6th International Conference on Learning Representations, ICLR 2018, Vancouver, BC, Canada, April 30 - May 3, 2018, Conference Track Proceedings. OpenReview.net, 2018. URL https://openreview.net/forum?id=B1l8BtlCb.

He, L., Zhang, S., Wu, L., Xia, H., Ju, F., Zhang, H., Liu, S., Xia, Y., Zhu, J., Deng, P., et al. Pre-training co-evolutionary protein representation via a pairwise masked language model. arXiv preprint arXiv:2110.15527, 2021.

Ho, J., Jain, A., and Abbeel, P. Denoising diffusion proba-bilistic models. Advances in Neural Information Processing Systems, 33:6840–6851, 2020.

Houlsby, N., Giurgiu, A., Jastrzebski, S., Morrone, B., De Laroussilhe, Q., Gesmundo, A., Attariyan, M., and Gelly, S. Parameter-efficient transfer learning for nlp. In International Conference on Machine Learning, pp. 2790–2799. PMLR, 2019.

Hsu, C., Verkuil, R., Liu, J., Lin, Z., Hie, B., Sercu, T., Lerer, A., and Rives, A. Learning inverse folding from millions of predicted structures. In Chaudhuri, K., Jegelka, S., Song, L., Szepesvari, C., Niu, G., and Sabato, S. (eds.), Proceedings of the 39th International Conference on Ma-chine Learning, volume 162 of Proceedings of Machine Learning Research, pp. 8946–8970. PMLR, 17–23 Jul 2022. URL https://proceedings.mlr.press/v162/hsu22a.html.

Hu, M., Yuan, F., Yang, K. K., Ju, F., Su, J., Wang, H., Yang, F., and Ding, Q. Exploring evolution-aware &-free protein language models as protein function predictors. In Advances in Neural Information Processing Systems, 2022.

Huang, P.-S., Boyken, S. E., and Baker, D. The coming of age of de novo protein design. Nature, 537(7620): 320–327, 2016.

Ingraham, J., Garg, V., Barzilay, R., and Jaakkola, T. Gener-ative models for graph-based protein design. In Advances in neural information processing systems, 2019.

Jin, W., Wohlwend, J., Barzilay, R., and Jaakkola, T. S. Iterative refinement graph neural network for antibody sequence-structure co-design. In International Confer-ence on Learning Representations, 2021.

Jing, B., Eismann, S., Suriana, P., Townshend, R. J. L., and Dror, R. Learning from protein structure with geomet-ric vector perceptrons. In International Conference on Learning Representations, 2020.

Jumper, J., Evans, R., Pritzel, A., Green, T., Figurnov, M., Ronneberger, O., Tunyasuvunakool, K., Bates, R., Žídek, A., Potapenko, A., et al. Highly accurate protein structure prediction with alphafold. Nature, 596(7873):583–589, 2021.

Kabsch, W. and Sander, C. Dictionary of protein secondary structure: pattern recognition of hydrogen-bonded and geometrical features. Biopolymers: Original Research on Biomolecules, 22(12):2577–2637, 1983.

Kaplan, J., McCandlish, S., Henighan, T., Brown, T. B., Chess, B., Child, R., Gray, S., Radford, A., Wu, J., and Amodei, D. Scaling laws for neural language models. arXiv preprint arXiv:2001.08361, 2020.

Kingma, D. P. and Ba, J. Adam: A method for stochas-tic optimization. In Bengio, Y. and LeCun, Y. (eds.), 3rd International Conference on Learning Represen-tations, ICLR 2015, San Diego, CA, USA, May 7-9, 2015, Conference Track Proceedings, 2015. URL http://arxiv.org/abs/1412.6980.

Kingma, D. P. and Welling, M. Auto-encoding variational bayes. In Bengio, Y. and LeCun, Y. (eds.), 2nd Interna-tional Conference on Learning Representations, ICLR 2014, Banff, AB, Canada, April 14-16, 2014, Conference Track Proceedings, 2014. URL http://arxiv.org/abs/1312.6114.

Kong, X., Huang, W., and Liu, Y. Conditional antibody design as 3d equivariant graph translation. arXiv preprint arXiv:2208.06073, 2022.

Lee, J., Tang, R., and Lin, J. What would elsa do? freezing layers during transformer fine-tuning. arXiv preprint arXiv:1911.03090, 2019.

Lees, J., Yeats, C., Perkins, J., Sillitoe, I., Rentzsch, R., Dessailly, B. H., and Orengo, C. Gene3D: a domain-based resource for comparative genomics, functional annotation and protein network analysis. Nucleic Acids Research, 40(D1):D465–D471, 12 2011. ISSN 0305-1048. doi: 10.1093/nar/gkr1181. URL https://doi.org/10.1093/nar/gkr1181.

Lefranc, M.-P., Pommié, C., Ruiz, M., Giudicelli, V., Foulquier, E., Truong, L., Thouvenin-Contet, V., and Lefranc, G. Imgt unique numbering for immunoglobulin and t cell receptor variable domains and ig superfamily v-like domains. Developmental & Comparative Immunol-ogy, 27(1):55–77, 2003.

Li, X. L. and Liang, P. Prefix-tuning: Optimizing continuous prompts for generation. In Proceedings of the 59th Annual Meeting of the Association for Computational Lin-guistics and the 11th International Joint Conference on Natural Language Processing (Volume 1: Long Papers), pp. 4582–4597, 2021.

Li, Z., Yang, Y., Faraggi, E., Zhan, J., and Zhou, Y. Direct prediction of profiles of sequences compatible with a protein structure by neural networks with fragment-based local and energy-based nonlocal profiles. Pro-teins: Structure, Function, and Bioinformatics, 82(10): 2565–2573, 2014. doi: https://doi.org/10.1002/prot.24620. URL https://onlinelibrary.wiley.com/doi/abs/10.1002/prot.24620.

Lin, Z., Akin, H., Rao, R., Hie, B., Zhu, Z., Lu, W., dos Santos Costa, A., Fazel-Zarandi, M., Sercu, T., Candido, S., et al. Language models of protein sequences at the scale of evolution enable accurate structure prediction. BioRxiv, 2022.

Lu, A. X., Zhang, H., Ghassemi, M., and Moses, A. Selfsupervised contrastive learning of protein representations by mutual information maximization. BioRxiv, pp. 2020–09, 2020.

Madani, A., Krause, B., Greene, E. R., Subramanian, S., Mohr, B. P., Holton, J. M., Olmos Jr, J. L., Xiong, C., Sun, Z. Z., Socher, R., et al. Deep neural language modeling enables functional protein generation across families. bioRxiv, pp. 2021–07, 2021.

McDermott, M., Yap, B., Hsu, H., Jin, D., and Szolovits, P. Adversarial contrastive pre-training for protein sequences. arXiv preprint arXiv:2102.00466, 2021.

Meier, J., Rao, R., Verkuil, R., Liu, J., Sercu, T., and Rives, A. Language models enable zero-shot prediction of the effects of mutations on protein function. In Advances in Neural Information Processing Systems, pp. 29287–29303, 2021.

Melnyk, I., Chenthamarakshan, V., Chen, P.-Y., Das, P., Dhurandhar, A., Padhi, I., and Das, D. Reprogramming large pretrained language models for antibody sequence infilling. arXiv preprint arXiv:2210.07144, 2022a.

Melnyk, I., Lozano, A., Das, P., and Chenthamarakshan, V. Alphafold distillation for improved inverse protein folding. arXiv preprint arXiv:2210.03488, 2022b.

Min, S., Park, S., Kim, S., Choi, H.-S., Lee, B., and Yoon, S. Pre-training of deep bidirectional protein sequence representations with structural information. IEEE Access, 9:123912–123926, 2021.

Nambiar, A., Heflin, M., Liu, S., Maslov, S., Hopkins, M., and Ritz, A. Transforming the language of life: transformer neural networks for protein prediction tasks. In Proceedings of the 11th ACM international conference on bioinformatics, computational biology and health informatics, pp. 1–8, 2020.

Nourani, E., Asgari, E., McHardy, A. C., and Mofrad, M. R. Tripletprot: deep representation learning of proteins based on siamese networks. IEEE/ACM Transactions on Computational Biology and Bioinformatics, 19 (6):3744–3753, 2021.

Olsen, T. H., Moal, I. H., and Deane, C. M. Ablang: An anti-body language model for completing antibody sequences. bioRxiv, 2022.

Orengo, C. A., Michie, A. D., Jones, S., Jones, D. T., Swindells, M. B., and Thornton, J. M. Cath–a hierar-chic classification of protein domain structures. Structure, 5(8):1093–1109, 1997.

Pfeiffer, J., Kamath, A., Rücklé, A., Cho, K., and Gurevych, I. Adapterfusion: Non-destructive task composition for transfer learning. arXiv preprint arXiv:2005.00247, 2020a.

Pfeiffer, J., Rücklé, A., Poth, C., Kamath, A., Vulić, I., Ruder, S., Cho, K., and Gurevych, I. Adapterhub: A framework for adapting transformers. arXiv preprint arXiv:2007.07779, 2020b.

Pfeiffer, J., Vulić, I., Gurevych, I., and Ruder, S. Mad-x: An adapter-based framework for multi-task cross-lingual transfer. arXiv preprint arXiv:2005.00052, 2020c.

Potter, S. C., Luciani, A., Eddy, S. R., Park, Y., Lopez, R., and Finn, R. D. HMMER web server: 2018 up-date. Nucleic Acids Research, 46(W1):W200–W204, 06 2018. ISSN 0305-1048. doi: 10.1093/nar/gky448. URL https://doi.org/10.1093/nar/gky448.

Qi, Y. and Zhang, J. Z. Densecpd: improving the accuracy of neural-network-based computational protein sequence design with densenet. Journal of chemical information and modeling, 60(3):1245–1252, 2020.

Radford, A., Narasimhan, K., Salimans, T., Sutskever, I., et al. Improving language understanding by generative pre-training. 2018.

Rao, R., Bhattacharya, N., Thomas, N., Duan, Y., Chen, P., Canny, J., Abbeel, P., and Song, Y. Evaluating protein transfer learning with tape. Advances in neural information processing systems, 32, 2019.

Rives, A., Meier, J., Sercu, T., Goyal, S., Lin, Z., Liu, J., Guo, D., Ott, M., Zitnick, C. L., Ma, J., and Fergus, R. Bi-ological structure and function emerge from scaling unsupervised learning to 250 million protein sequences. PNAS, 2019. doi: 10.1101/622803. URL https://www.biorxiv.org/content/10.1101/622803v4.

Rombach, R., Blattmann, A., Lorenz, D., Esser, P., and Ommer, B. High-resolution image synthesis with latent diffusion models. In Proceedings of the IEEE/CVF Con-ference on Computer Vision and Pattern Recognition, pp. 10684–10695, 2022.

Salazar, J., Liang, D., Nguyen, T. Q., and Kirchhoff, K. Masked language model scoring. In Proceedings of the 58th Annual Meeting of the Association for Computa-tional Linguistics, pp. 2699–2712, 2020.

Savinov, N., Chung, J., Binkowski, M., Elsen, E., and van den Oord, A. Step-unrolled denoising autoencoders for text generation. In International Conference on Learning Representations, 2021.

Sennrich, R., Haddow, B., and Birch, A. Improving neural machine translation models with monolingual data. In Proceedings of the 54th Annual Meeting of the Associ-ation for Computational Linguistics (Volume 1: Long Papers), pp. 86–96, Berlin, Germany, August 2016. As-sociation for Computational Linguistics. doi: 10.18653/v1/P16-1009. URL https://www.aclweb.org/anthology/P16-1009.

Sohl-Dickstein, J., Weiss, E., Maheswaranathan, N., and Ganguli, S. Deep unsupervised learning using nonequi-librium thermodynamics. In International Conference on Machine Learning, pp. 2256–2265. PMLR, 2015.

Steinegger, M. and Sö ding, J. Mmseqs2 enables sensi-tive protein sequence searching for the analysis of mas-sive data sets. Nature biotechnology, 35(11):1026–1028, 2017.

Street, A. G. and Mayo, S. L. Computational protein design. Structure, 7(5):R105–R109, 1999.

Strodthoff, N., Wagner, P., Wenzel, M., and Samek, W. Udsmprot: universal deep sequence models for protein classification. Bioinformatics, 36(8):2401–2409, 2020.

Sturmfels, P., Vig, J., Madani, A., and Rajani, N. F. Profile prediction: An alignment-based pre-training task for pro-tein sequence models. arXiv preprint arXiv:2012.00195, 2020.

Su, J., Lu, Y., Pan, S., Murtadha, A., Wen, B., and Liu, Y. Roformer: Enhanced transformer with rotary position embedding. arXiv preprint arXiv:2104.09864, 2021.

Suzek, B. E., Wang, Y., Huang, H., McGarvey, P. B., Wu, C. H., and Consortium, U. Uniref clusters: a comprehen-sive and scalable alternative for improving sequence sim-ilarity searches. Bioinformatics, 31(6):926–932, 2015.

Tan, C., Gao, Z., Xia, J., and Li, S. Z. Generative de novo protein design with global context. arXiv preprint arXiv:2204.10673, 2022.

Towse, C.-L. and Daggett, V. When a domain isn’ta domain, and why it’s important to properly filter proteins in databases: Conflicting definitions and fold classification systems for structural domains makes filtering of such databases imperative. BioEssays: news and reviews in molecular, cellular and developmental biology, 34(12): 1060, 2012.

Unsal, S., Atas, H., Albayrak, M., Turhan, K., Acar, A. C., and Doğan, T. Learning functional properties of proteins with language models. Nature Machine Intelligence, 4 (3):227–245, 2022.

van den Burg, G. and Williams, C. On memorization in probabilistic deep generative models. Advances in Neural Information Processing Systems, 34:27916–27928, 2021.

Vaswani, A., Shazeer, N., Parmar, N., Uszkoreit, J., Jones, L., Gomez, A. N., Kaiser, L., and Polosukhin, I. Attention is all you need. In Guyon, I., von Luxburg, U., Bengio, S., Wallach, H. M., Fergus, R., Vishwanathan, S. V. N., and Garnett, R. (eds.), Advances in Neural Information Processing Systems 30: Annual Conference on Neural Information Processing Systems 2017, December 4-9, 2017, Long Beach, CA, USA, pp. 5998–6008, 2017. URL https://proceedings.neurips.cc/paper/2017/hash/3f5ee243547dee91fbd053c1c4a845aa-Abstract.html.

Verkuil, R., Kabeli, O., Du, Y., Wicky, B. I., Milles, L. F., Dauparas, J., Baker, D., Ovchinnikov, S., Sercu, T., and Rives, A. Language models generalize beyond natural proteins. bioRxiv, pp. 2022–12, 2022.

Watson, J. L., Juergens, D., Bennett, N. R., Trippe, B. L., Yim, J., Eisenach, H. E., Ahern, W., Borst, A. J., Ragotte, R. J., Milles, L. F., et al. Broadly applicable and accurate protein design by integrating structure prediction networks and diffusion generative models. bioRxiv, pp. 2022–12, 2022.

Wei, J., Tay, Y., Bommasani, R., Raffel, C., Zoph, B., Borgeaud, S., Yogatama, D., Bosma, M., Zhou, D., Metzler, D., et al. Emergent abilities of large language models. Transactions on Machine Learning Research, 2022a.

Wei, J., Wang, X., Schuurmans, D., Bosma, M., Xia, F., Chi, E. H., Le, Q. V., Zhou, D., et al. Chain-of-thought prompting elicits reasoning in large language models. In Advances in Neural Information Processing Systems, 2022b.

Xiao, Y., Qiu, J., Li, Z., Hsieh, C.-Y., and Tang, J. Modeling protein using large-scale pretrain language model. arXiv preprint arXiv:2108.07435, 2021.

Yang, K. K., Wu, Z., and Arnold, F. H. Machine-learningguided directed evolution for protein engineering. Nature methods, 16(8):687–694, 2019.

Yang, K. K., Lu, A. X., and Fusi, N. Convolutions are competitive with transformers for protein sequence pre-training. bioRxiv, pp. 2022–05, 2022a.

Yang, K. K., Zanichelli, N., and Yeh, H. Masked inverse folding with sequence transfer for protein representation learning. bioRxiv, pp. 2022–05, 2022b.

Zhu, J., Shang, Y., Xia, C., Wang, W., Wen, W., and Zhang, M. Guanylate kinase domains of the maguk family scaf-fold proteins as specific phospho-protein-binding mod-ules. The EMBO journal, 30(24):4986–4997, 2011.

